# Embryo-scale single-cell chemical transcriptomics reveals dependencies between cell types and signaling pathways

**DOI:** 10.1101/2025.04.03.646423

**Authors:** Eliza Barkan, Madeleine Duran, Nicholas Lammers, Amy Tresenrider, Dana Jackson, Heidi Lee, Benjamin Haagen, Lauren Saunders, Phil Abitua, David Kimelman, Cole Trapnell

## Abstract

Organogenesis is a highly organized process that is conserved across vertebrates and is heavily dependent on intercellular signaling to achieve cell type identity. We lack a comprehensive understanding of how developing cell types in each organ and tissue depend on developmental signaling pathways. To address this gap in knowledge, we captured the molecular consequences of inhibiting each of the seven major developmental signaling pathways in zebrafish, using large-scale whole embryo single cell RNA-seq from over two million cells. This approach allowed us to detect signaling pathway regulation even in very rare cell types. By focusing on the development of the pectoral fin, we uncovered two new cell types (distal mesenchyme and tenocytes) and multiple novel signaling dependencies during pectoral fin development. This resource serves as a valuable tool for investigators seeking to rapidly assess the role of the major signaling pathways during the formation of their tissue of interest.

## Introduction

Intercellular signaling is central to embryonic development. Gradients of secreted factors such as Wnt, Bone Morphogenetic Protein (BMP), Fibroblast Growth Factor (FGF) and others pattern the early embryo, drive gastrulation, and direct tissue morphogenesis. These pathways are critical for coordinating fate decisions of cells from different lineages, controlling their proliferation, migration, sorting, and apoptosis^1,2^. Despite their central role in diverse tissues and organs, studying how signaling pathways direct embryogenesis has been difficult, particularly in vertebrates. One major challenge is the extensive redundancy of receptors and ligands within vertebrate genomes^2^. Mutants for individual ligands or receptors in a particular signaling pathway often have mild phenotypes, typically because of redundant signals or because cells lacking one receptor can still receive signals through another from the same family^3^. Secondly, signals are used repeatedly during development, but their outcomes are highly time-dependent. Disrupting a pathway at the start of development can have very different effects compared to altering it at a later time. A third concern for many organisms such as fish and frogs, which have large maternal expression of signaling components, is that a mutant (or crispant) that alters a key component of a pathway may not show a phenotype until much later in development, if at all, because the maternal contribution is sufficient to support early development. Finally, and most importantly, our knowledge of the effects of signaling on specific tissues and organs is primarily derived from selective perturbations and phenotypic analyses that interrogate effects on one or just a few cell lineages that were of interest to a specific investigator. For all of these reasons, we lack a comprehensive understanding of the role of intercellular signaling in early vertebrate development.

In this work, we leverage recent advances in sequencing technology to overcome key challenges of studying signaling dependencies during organogenesis. Improvements to the throughput of single cell genomics have enabled the measurement of whole transcriptomes from many individually barcoded embryos^4^. When applied to embryos that have been perturbed with genome editing^4^ or through environmental stress^5^, these protocols have proven fruitful for defining the genetic requirements of individual cell types and the molecular mechanisms that give rise to underlying anatomic phenotypes. In this study, we leveraged the external development, small size and high fecundity of zebrafish to enable precise perturbation of signaling pathways through the addition of well-characterized small molecule inhibitors directly to the surrounding medium, which allows us to overcome issues of redundant receptors and ligands within each signaling pathway, and allows us to examine the effects of pathway inhibition at different developmental stages. By profiling zebrafish embryos treated with inhibitors for the seven major signaling pathways of early development, we sought to define the dependencies of each cell type on these pathways during organogenesis in a single high-throughput experiment.

Here, we report CHEMFISH, an atlas of over two million single-cell transcriptomes from more than 500 individual zebrafish embryos in which signaling pathways have been disrupted using small molecules. By adding inhibitors against BMP, FGF, Notch, Retinoic acid (RA), Hedgehog (Hh), Transforming Growth Factor-β (TGF-β), and Wnt, we define how the abundance and transcriptomes of cell types in the developing embryo require each pathway. Adding inhibitors at five timepoints, from the start of gastrulation through the pharyngula stage (6 (shield), 13 (8-somite), 24, 36 and 42 hours post fertilization (hpf)^6^), enabled us to distinguish direct effects from earlier effects that propagate through the lineage (**Figure 1A**). Although these perturbations impact nearly every cell type in the embryo, we examine the zebrafish pectoral fin as an in-depth example to demonstrate the value of our approach for discovering new cell types and new roles for pathways in even well-studied organs. For example, we found a previously uncharacterized role for TGF-β in the differentiation of all the lateral plate mesoderm (LPM)-derived pectoral fin cell types, as well as a surprising inhibitory role of Hh in regulating the development of the pectoral fin support bone (cleithrum). Transcriptional analysis, which we validate through a new imaging-based 3D model of the fin, characterized a recently reported but incompletely understood tendon-forming cell (tenocyte) population in between the muscle and core cartilage that we show differentiates from the initial pectoral fin progenitors. We show that another cell type, the ‘distal mesenchyme,’ previously described as originating from the apical ectodermal ridge (AER)^7,8^, also differentiates from this same progenitor population. Finally, we find that BMP negatively regulates AER and distal mesenchyme, and its inhibition results in aberrant blood vessel distribution in the pectoral fin. Our results demonstrate this new extensive database will be broadly useful for modeling tissue-specific signaling pathway regulation of early vertebrate development at embryo-scale.

**Figure 1.**
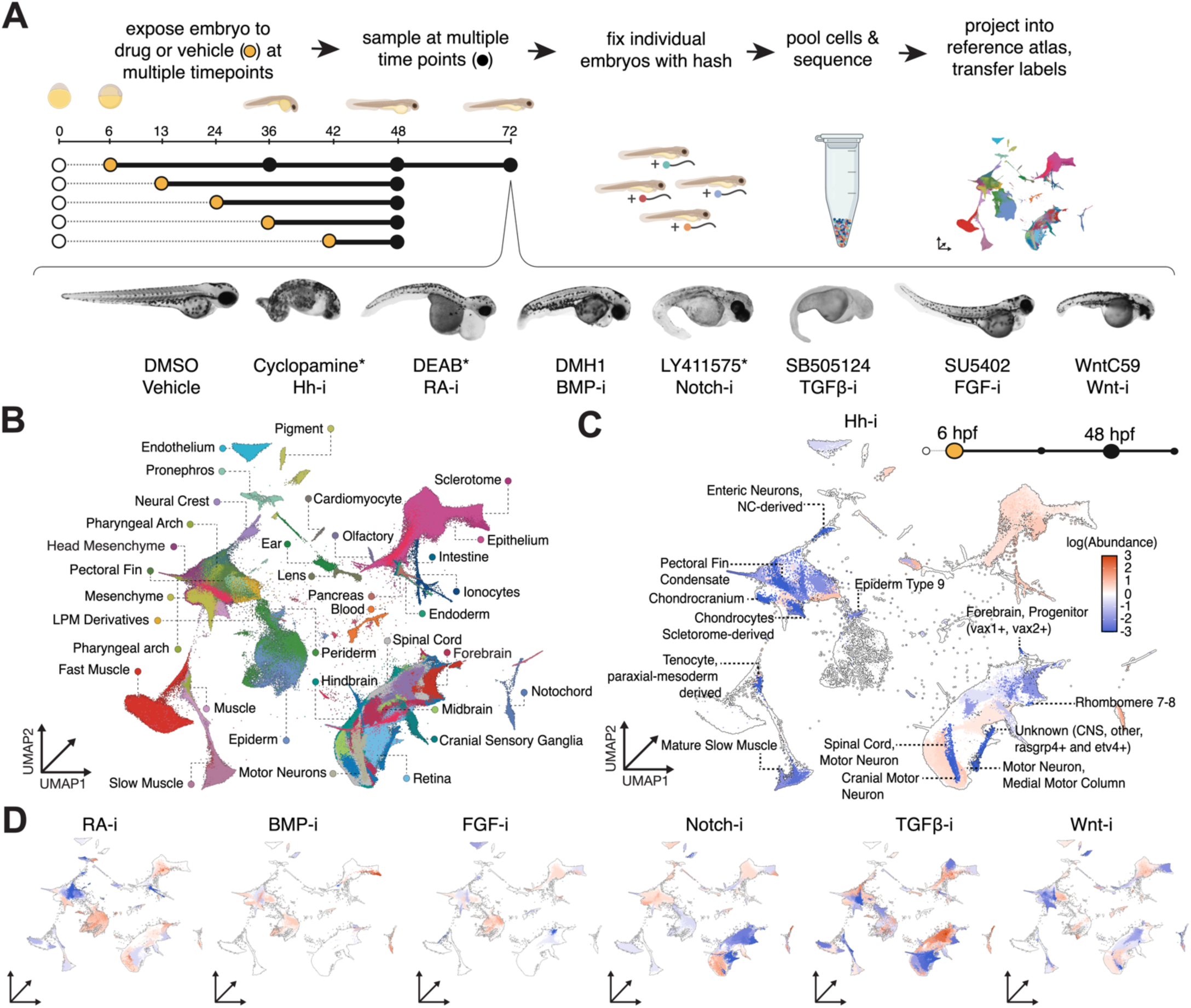
Effects of signaling pathway inhibitors on organogenesis are captured with individual embryo hashing of single-cell transcriptomes. A) Experimental workflow of signaling pathway perturbations and individual embryo hashing. Representative images of 72 hpf embryos raised in vehicle (DMSO) or inhibitor. * indicates select perturbations with data from a 6 hpf inhibitor addition. B) Chemically perturbed transcriptional data projected into reference atlas UMAP and colored and labeled by tissue. C) UMAP embedding, colored by significant (q<0.05, Poisson log-normal (PLN) regression) log fold change (logFC) in abundance of each cell type in cyclopamine-treated, Hh-i embryos. Cell types labeled in example Hh-i have absolute value of abundance logFC > 3 (n=8). D) UMAP embeddings, colored by significant (q<0.05, PLN regression) logFC in abundance of each cell type for each inhibitor added at the earliest time point (6 or 13 hpf) and collected at 48 hpf, compared to matched vehicle control (n=8).

## Results

### Multiplexed single cell RNA-seq profiles signaling pathway regulation of organogenesis

To detect effects of signaling pathway inhibitors on changes to cellular abundance and gene expression, we used sci-Plex to capture single nucleus transcriptomes from hundreds of individual zebrafish embryos^4,9^. We designed these experiments similarly to our previous genetic^4^ and environmental perturbation studies^5^, to profile thousands of nuclei from each embryo. Each embryo analyzed was individually tagged with a unique oligonucleotide (a barcode sequence called a ‘hash’) in order to uniquely track each perturbation, time point at which the inhibitor was added, and the time point the embryo was collected, which allowed us to confer sufficient statistical power to infer how each perturbation impacts even very rare cell types (**Figure 1A**, **Figure S1A-F**) (Methods).

In the first round of experiments, chemical inhibitors of the major developmental signaling pathways (BMP, FGF, Notch, RA, Hh, TGF-β & Wnt) were added to zebrafish embryos. Because inhibition of many of these signaling pathways (BMP, FGF, TGF-β & Wnt) cause such severe effects during gastrulation^10,11^, we waited to inhibit embryos until the early somite stages (13 hpf). However, since inhibiting a few of the pathways (Notch, RA, Hh) did not cause major defects during gastrulation, we added these three inhibitors at 6 hpf to be able to fully capture their roles during development. To disentangle direct vs indirect effects, we also added the inhibitors at multiple time points during organogenesis (13, 24, 36, 42 hpf) in a second round of experiments.

To label the cells in our experiment by cell type, we projected our cells into our developmental zebrafish atlas comprising 1.2 million cells, spanning time points from 18 hpf to 96 hpf, and transferred major tissue-level (30) and cell type labels (319) to each cell in the experiment from its nearest neighbors in the reference (**Figure 1B**)^4,12^ (Methods) (Duran *et al.,* see related manuscript file). Cells of many different types were captured from each embryo, with an average of 2,684 total cells captured per embryo (**Figure S1E,G-I**). This database provides a strong foundation for examining signaling dependent cell type abundance changes during vertebrate organogenesis.

### Differential cell type abundance reveals diverse utilization of all signaling pathways in organogenesis

In order to define the cell types that require each signaling pathway, we compared inhibitor-treated embryos to their vehicle-treated controls using the newly developed R package Hooke^12^ (Duran *et al*., see related manuscript file). Briefly, Hooke models how cell type proportions change over time and in response to perturbations using a Poisson log-normal (PLN) Network^13^. This comparison detected numerous cell types that were significantly depleted (median = 83) or enriched (median = 76) relative to matched controls, across each drug addition time point and collection time point (FDR < 5%; see Methods).

Each perturbation had a unique and broad pattern of differential cell type abundances across the different tissues and cell types in the embryo (**Figure 1C-D**; controls shown in **Figure S2A**). As expected, UMAP embedding of the individual cell type counts per embryo demonstrated that individual embryos clustered with other embryos treated with the same chemical inhibitor (**Figure 2A**). Moreover, when we examined two different inhibitors that have previously been shown to inhibit the TGF-β pathway^14,15^, we found that these embryos clustered together in UMAP space due to similar differential cell type abundance (**Figure 2A**, **Figure S2B-C**), reaffirming the specificity of the inhibitor treatments. Additionally, inhibiting each pathway altered the relative abundances of cell types in a distinct way. For example, consistent with the literature^16–19^, early treatment (13 hpf) with inhibitors of FGF, Wnt and RA signaling led to dramatic reductions in pectoral fin abundance at 48 hpf, whereas unlike RA inhibition, inhibition of FGF and Wnt starting at 24 hpf (11 hours later) continued to impact the abundance of pectoral fin mesoderm (**Figure 2B**), demonstrating that our approach can distinguish cell-type specific signaling requirements in the early embryo from dependencies that manifest later in organogenesis.

**Figure 2.**
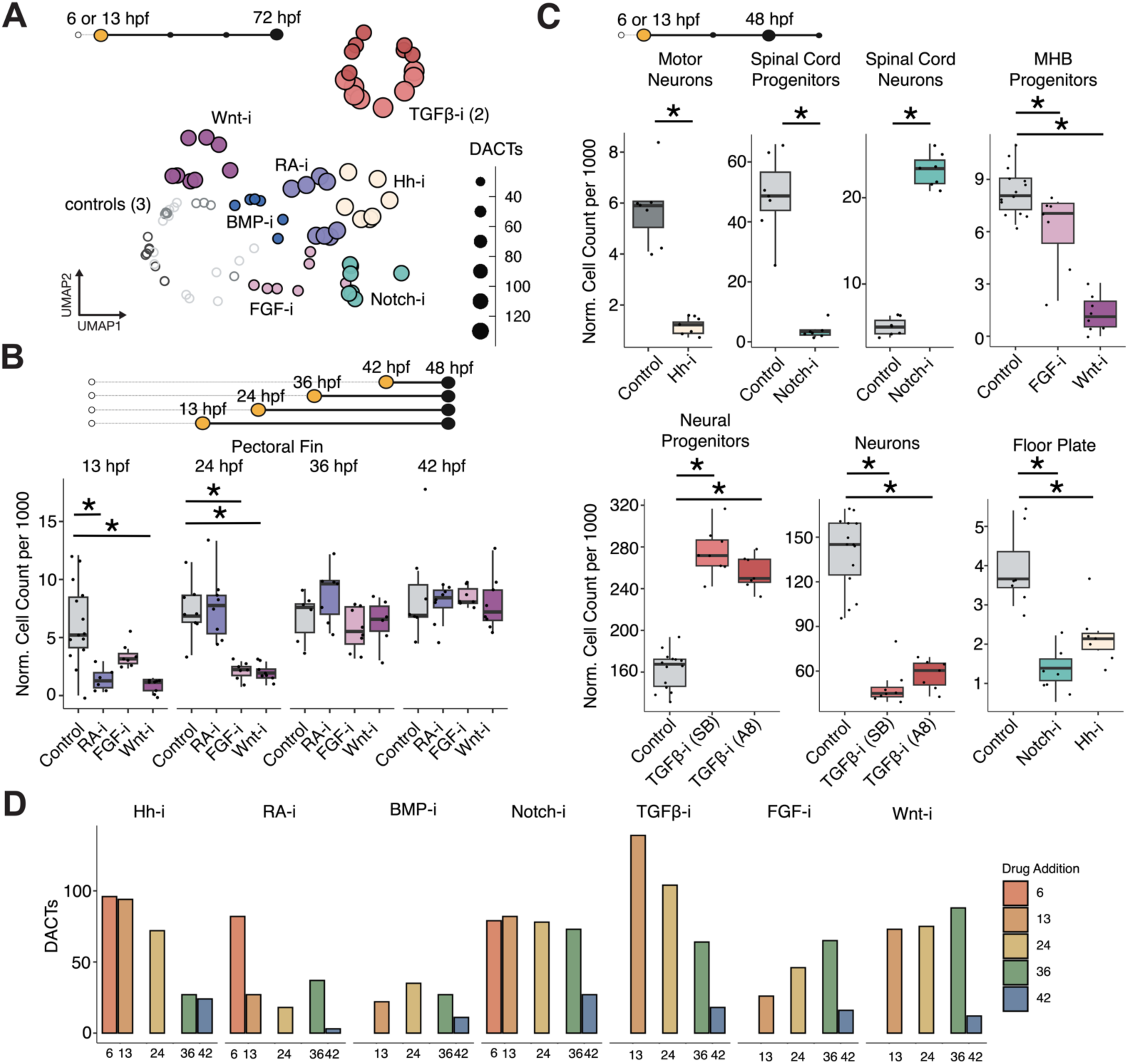
Whole embryo high-resolution abundance phenotyping of chemically inhibited embryos. A) UMAP embedding of the individual cell type counts per embryo at 72 hpf. Each dot is an embryo colored by inhibitor and sized by the number of differentially abundant cell types (DACTs) when compared to matched vehicle controls, DMSO (light gray), EtOH (medium gray) or no vehicle (dark gray). In the case of the TGF-β pathway inhibitors that were examined: A8301 is colored dark red and SB505124 is colored light red. B) Size-factor normalized cell count per 1000 cells in each embryo for select inhibitors and matched control for the pectoral fin mesoderm tissue. Significance (*q<0.05, PLN regression) relative to matched vehicle control embryos. C) Size-factor normalized cell count per 1000 cells in each embryo for each control and inhibitor for each broad cell type. Broad cell types are pooled from multiple finer cell type annotations. MHB, midbrain-hindbrain boundary. Spinal cord progenitors and spinal cord differentiated neurons, floor plate and motor neuron boxplots are from 36 hpf, all other data shown in boxplots are from 48 hpf. Significance (*q<0.05, PLN regression) relative to matched vehicle control embryos. D) Bar plot of the number of DACTs per inhibitor addition and perturbation for the 48 hpf collection time point.

We next sought to confirm that our inhibitor treatments recapitulated the known signaling requirements in various cell types throughout the embryo, starting with the central nervous system (CNS) because it is well-documented. For example, in our own data we recapitulated that Shh positively regulates the formation of motor neurons^20,21^, Notch negatively regulates the differentiation of spinal cord neurons^22^, FGF and Wnt positively regulate the formation of the midbrain-hindbrain boundary^23^, TGF-β positively regulates neuron differentiation in many CNS cell types^24,25^ and Hh and Notch positively regulate formation of the floor plate^26,27^ (**Figure 2C**, **Figure S2D**). Outside of the nervous system, we recapitulated numerous other instances where signaling pathways regulated the abundance of a cell type, for example Shh positively regulates slow muscle^28,29^ and formation of pectoral fin cartilage condensate^30,31^ (**Figure 1C**). During zebrafish pronephros development, RA is necessary for specifying proximal tubule fates and loss of RA causes an expansion of distal tubule segments^32^. Consistent with this role of RA, we observed that RA-inhibition beginning at shield stage led to a loss of three proximal cell types and a dramatic increase in the abundance of two distal cell types in the pronephros at 48 hpf (**Figure S2E**).

Having confirmed that our sci-Plex experiments captured the expected signaling requirements of cell types on signaling pathways, we turned to enriched or depleted cell types that were not already known to depend on the pathways we targeted. For example, when we measured the effects of two TGF-β inhibitors, SB505124 and A8301, our high-resolution abundance phenotyping identified many diverse roles of TGF-β in organogenesis that are shared across both inhibitors (**Figure 1D**, **Figure S2B-C,D**). For example, we observed a complete loss of alcian blue stained cartilage in the head and pectoral fins of TGFβ-i embryos (**Figure S2F**). Some cell types, such as neural progenitors, parachordal cartilage and mesenchyme-fated neural crest, were significantly depleted in embryos in which TGF-β was added at 13 hours, despite the absence of obvious anatomic phenotypes (**Table S1**). Taken together, the combination of the sequencing data with the Hooke analysis package, allows the rapid identification of specific signaling-dependent changes in hundreds of cell types, as well as the ability to detect subtle changes in individual cell types or tissues that would be overlooked with standard phenotyping that assess macroscopic features or tissue morphology.

Of the embryos collected at the pharyngula stage (48 hpf), embryos exposed to pathway inhibitors at earlier time points showed effects in more cell types than embryos treated later in development. The mean number of differentially abundant cell types (DACTs) per inhibitor addition was consistent with temporally specific roles of signaling in patterning the embryo after gastrulation (**Figure 2D**). Notably, later inhibition of Notch, Wnt, FGF and TGF-β continued to yield many DACTs. In contrast, RA tended to be selectively important early on (**Figure 2D**), exemplified by RA’s specific early regulation of pectoral fin development (**Figure 2B**).

### Identification of molecular targets of signaling pathways

We next asked how the transcriptome of each cell type was impacted by the chemical perturbations. Briefly, we compared each gene’s expression in chemically inhibited embryos by first summing gene expression of each cell by their cell type and embryo of origin, yielding 18,216 total pseudobulks. We then fit a quasipoisson-valued generalized linear model to each gene’s pseudobulked values, weighting pseudobulks by the number of cells within it (Methods). These models estimated the effect of an inhibitor on each gene’s expression in each cell type. However, because the inhibitors also altered cell type proportions throughout the embryo, we limited the comparison to embryos treated with inhibitors at 42 hpf, six hours prior to collection at 48 hpf. While few cell types were differentially abundant in this contrast (**Figure 2D**), the analysis nevertheless detected many differentially expressed genes (DEGs) per perturbation (median = 2229, interquartile range (IQR) = 1935-2393) when pooled across all cell types. DEGs were more often found in abundant cell types than rare ones, a detection bias expected for count-based regression methods for differential expression. Nevertheless, we captured pathway-specific effects within rare cell types such as in pectoral fin and cranial sensory ganglia (**Figure 3A**, **Figure S3A,B**).

**Figure 3.**
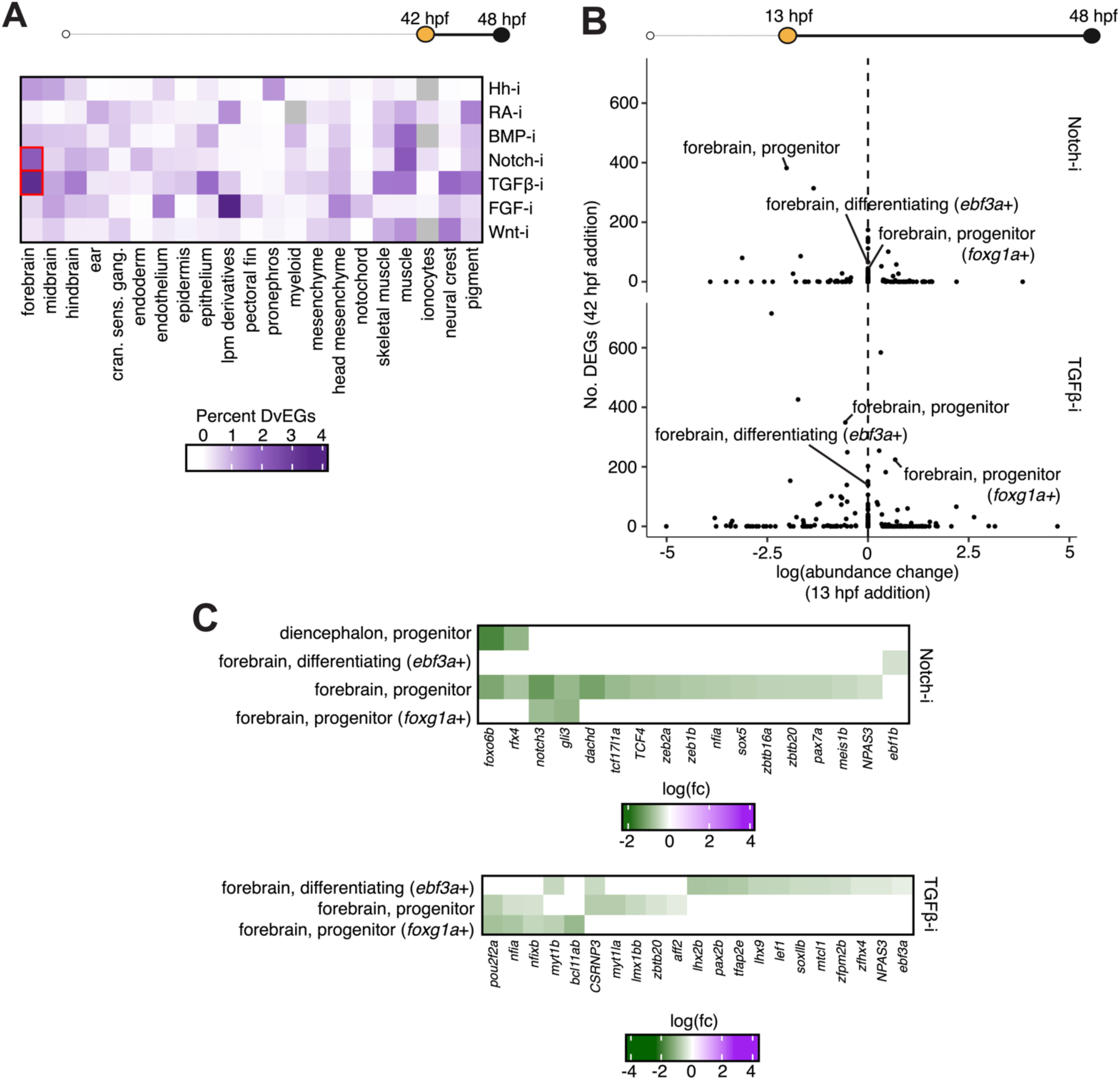
Systematic detection of cell-type specific deviantly expressed genes. A) Heatmap of the log10 percent of differentially expressed genes that change as cells transition from one state to another in wild type embryos (‘DvEGs’), as defined by Platt (q<0.05) in each tissue, that are also DEGs for the 42 hpf addition of all inhibitors, collected at 48 hpf. B) Scatterplot of the abundance of each cell type in LY411575 treated, Notch-i and SB505124 treated, TGFβ-i embryos compared to the number of DEGs for each cell type in each perturbation. Abundance changes plotted are from the 13 hpf inhibitor addition. C) Heatmap of the log10 fold change (FC) in gene expression for the DvEGs with q < 0.005 in forebrain cell types of LY411575 treated, Notch-i and SB505124 treated, TGFβ-i embryos.

To further validate our approach for interrogating the molecular targets of each signaling pathway, we demonstrate that we observe the expected changes in expression of known transcriptional targets of each signaling pathway (**Figure S3C**). For example, *gli1*, which operates downstream of Hh signaling^33^, was significantly downregulated in cell types of forebrain, spinal cord and connective tissue (**Figure S3C,D**).

Likewise, multiple Notch pathway targets including *dld, dtx4b*, *notch1b* and *notch3* themselves were down in Notch-i embryos (**Figure S3C**)^33^. However, not all canonical targets were significantly affected in cell types that express them. In Hh-i embryos, *gli1* changes were more likely to be found in abundant cell types that also express *gli1* at high levels than in rare cell types or those that express *gli1* at low levels (**Figure S3D,E**).

Moreover, we observed changes that might be due to cross-talk between the pathways: for example, *her6*, an important target and effector of Notch signaling^33^, was also downregulated in Hh-i embryos in many cell types (**Figure S3C**), including chondrocyte-fated neural crest, basal cells and other connective tissue. Thus, while our approach to transcriptionally phenotype at single-cell resolution can indeed detect the targets of the pathways we inhibited, deciphering the direct targets of each pathway is dependent on sampling depth of the gene, cell, and specimen levels.

Our sequencing data provided a rich resource for identifying novel downstream genes. We looked at the share of DEGs that were transcription factors (TFs). This analysis identified tens to hundreds (median = 189, IQR = 177-201) of TFs that were responsive to each signaling pathway, either globally or in a cell type specific manner (**Figure S3B,F**). To eliminate as many indirectly regulated genes as possible, we focused on DEGs that were concordantly affected in multiple cell types. Reassuringly, the 10 most shared differentially expressed transcription factors included numerous known targets. For example, *smad7* and *stat3*, both important effectors of TGF-β signaling^33^, were differentially expressed in numerous cell types in TGFβ-i embryos (**Figure S3F**). Together, these analyses demonstrate our atlas of chemically perturbed embryos can be used to identify targets of major signaling pathways.

Finally, we aimed to understand the molecular roles of each pathway in governing individual fate decisions within the embryo. To do so, we began with a list of genes that change as cells transition from one cell type to another in wild type embryos, as defined by Platt^12^ (Duran *et al.*, see related manuscript file). Briefly, Platt assembles a map of cell types transitions. Platt works by integrating single-cell transcriptome time series and perturbation experiments. We recently applied it to our atlas of wild-type and genetically perturbed embryos to construct a draft transition map of the zebrafish from 18 to 96 hpf ^4,12^ (Duran *et al.*, see related manuscript file).

We then used the map to define the genes that change as control cells progress through different stages of development^12^ (Duran *et al.*, see related manuscript file). The catalog of gene expression changes that occur as each cell type differentiates is useful for understanding how a perturbation disrupts each cell type’s development. Here, we tested whether the changes to each that occur in the reference atlas still occur as expected when specific signaling pathways are inhibited. We deem genes that fail to do so “deviantly expressed genes (DvEGs)” (Methods). For example, a gene that would normally be upregulated as a cell commits to a particular fate but fails to reach the same level in chemically-treated progenitors committing to a particular fate would be DvEG in that cell fate. Of the 587,177 genes that were upregulated or downregulated in control cells upon entry into a new cell state, 21,384 were DvEG when at least one pathway was inhibited. As expected, we capture many DvEGs in tissues that are known targets of signaling pathways, for example, FGF and RA in LPM derivatives^34,35^, FGF in pectoral fin^19^, Wnt in neural crest^36^, TGF-β and Notch in neural tissues^25,37^ and Hh in pronephric development^38,39^ (**Figure 3A**).

TGF-β and Notch have opposing roles in regulating neuronal differentiation^24,40^, and interestingly, we captured many DvEGs in both perturbations (**Figure 3A**). We noticed that although some forebrain progenitor cell types were not differentially abundant, we still captured many DvEGs in these cell types in both perturbations (**Figure 3B**). When we looked more closely, we noticed that some of the DvEGs included classic targets of the pathways discussed above. For example, *notch3*, normally upregulated in forebrain neural progenitors undergoing a transition to differentiated neurons of the zona limitans intrathalamica, failed to reach the expected levels of expression in Notch-i embryos. We also noticed targets of other pathways such as *gli3* (Hh), *tcf17l1a* (Wnt) and *TCF4* (Wnt) in Notch-i and *lef1* (Wnt) in TGFβ-i^33^, suggesting cross regulation of the Hh and Wnt pathways, which are also known regulators of CNS development^23,26,41^.

Numerous transcription factors normally upregulated in and required by specific cell types were aberrantly expressed in inhibitor-treated embryos. With TGF-β inhibition, we found *lhx2b* and *lhx9* failed to reach the expected levels of expression in differentiating forebrain cells. Lhx TFs are critical for forebrain differentiation^42^ but have no known dependence on TGF-β signaling. Additionally we found that *bcl11ab* and *myt1la*, which are known to be selectively expressed in two forebrain progenitor cell types, were depleted in TGF-β inhibition (**Figure 3C**). Mutations in these TFs cause reduced head and brain size in zebrafish and intellectual disability in humans yet also have no known dependence on TGF-β signaling^43,44^. Taken together these analyses demonstrate an example of how our data captured the molecular impact of disrupted signaling.

### A high-resolution transcriptional map of pectoral fin mesoderm

The main value to a comprehensive interrogation of the role of different signaling pathways is to guide research in new directions. We were struck looking at our data that every signaling pathway had effects on the formation of the pectoral fin. The pectoral fin is of particular interest as it is the antecedent to the tetrapod limb, with many of the fundamental developmental mechanisms being highly conserved^45^.

The zebrafish pectoral fin is made up of cell types arising from multiple origins: lateral plate mesoderm, somitic mesoderm (both sclerotome and myotome) and ectoderm. In the zebrafish, between 10 and 16 hours post fertilization (hpf), spatially segregated RA and Wnt signaling in the lateral plate mesoderm induces expression of the transcription factor *tbx5a*^16–18^. In the mesoderm, Tbx5a initiates a cascade of FGF expression^19,46^, which subsequently induces FGF expression in the adjacent ectoderm (AER) in the outer tip of the appendage^7,47,48^. Feed-forward loops maintain FGF expression as the appendage bud grows out^19^. Shh, known in amniotes for its role in forming and patterning digits^49^ is essential in zebrafish for maintaining and growing the paired appendage^30,31,50^, although the role of Shh in individual pectoral fin cell types is not known. After the fin bud has formed, BMP inhibits fin outgrowth by inhibiting *smoc1* expression^51^. Thus, the zebrafish pectoral fin requires signaling from Wnt, FGF, RA, Hh, and BMP, but how their molecular roles vary within individual fin cell types is not clear, and whether TGF-β and Notch play roles in pectoral fin formation is unknown.

On the mesoderm UMAP we identified a region that expresses *tbx5a*, and a neighboring region of cells that expresses high levels of *runx2b* (**Figure 4A**). Whereas *tbx5a* marks the pectoral fin progenitors, *runx2b* identifies the cells forming the cleithrum, which is an ossified structure that connects the pectoral fin to the cranium, analogous to the amniote shoulder blade^52^. We then reclustered this region of the mesoderm UMAP and observed multiple trajectories originating from the *tbx5a* progenitor population beginning at the earliest time point (**Figure 4B,C**). These progenitors, which also expressed *fgf24*, developed into a *fgf10a* mesodermal population (**Figure S4A**), consistent with the known role of these FGF signaling factors in regulating early pectoral fin development^19,48^. Contrasting cell type composition of the pectoral fin mesoderm from 20 somites (18 hpf) to larval stage (96 hpf), incrementally, *Platt*^12^ (Duran *et al.*, see related manuscript file) was used to define the lineage relationships of cell types in a directed graph. This revealed that the bud progenitors differentiate into the *fgf10a*+ fin bud mesoderm which, in turn, gives rise to four different cell types at 36-48 hpf (**Figure 4D**). Of these four cell types, two– the *sox9a*+ cartilage condensate and *runx2b*+ cleithrum– correspond to known populations (**Figure S4A**). The other two populations were not immediately identifiable.

**Figure 4.**
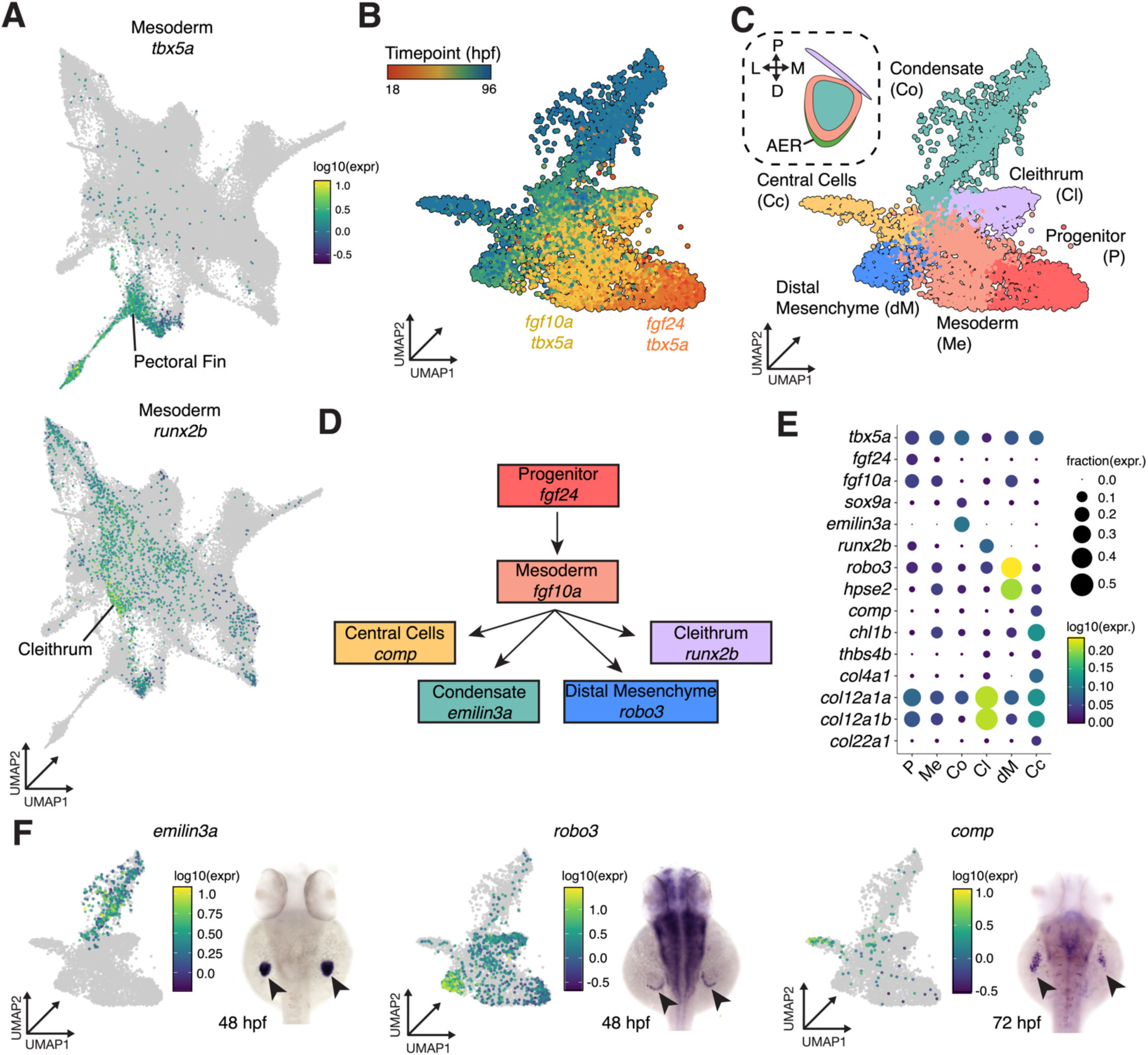
Trajectory and gene marker analysis of pectoral fin mesoderm. A) Sub-clustered UMAP of all mesoderm cells in our developing zebrafish atlas. The *tbx5a* gene broadly labels lateral plate mesoderm-derived pectoral fin mesoderm and *runx2b* marks the cleithrum (n = 15,793 cells). B) Sub-UMAP of *tbx5a*+ and *runx2b*+ cell populations colored by time point and C) our cell type annotations. Diagram in box represents known spatial representation of cell types at 48 hpf. D) Proposed pectoral fin mesoderm cell lineage diagram, derived from abundance of cells from control embryos over time, using *Platt* R package. E) Dot plot heatmap of selected canonical and novel top marker genes, grouped by cell type, colored by log10 gene expression, dots sized by the fraction of cells expressing the gene marker. F) UMAP expression plots (left) and whole mount *in situ* hybridization (WISH) (right) for three novel genes specific to pectoral fin cell types, *emilin3a* (condensate), *robo3* (distal mesenchyme), *comp* (central cells).

Next, we used DEG analysis to identify unique markers for each of the four terminal cell types found to make up the pectoral fin (**Table S2**, **Figure 4E**). We found that the *sox9a*+ cartilage condensate, which runs down the middle of the pectoral fin, is uniquely marked by the glycoprotein gene *emilin3a* (**Figure 4F**, left, **Figure S4A,B**). A second (unknown) cell type was marked by strong expression of the neural guidance gene *robo3* (**Figure 4F**, center). *robo3* was previously thought to be selectively expressed in the ectodermal cells that comprise the pectoral fin AER based on whole mount in situ hybridization analysis^53^, but our data clearly shows that *robo3* is expressed within a distinct mesoderm cell population (**Figure 4E**) and *not* in the AER (see below). We labeled these cells ‘distal mesenchyme.’ In addition to selectively expressing the axon guidance gene, *robo3*, the distal mesenchyme specifically expresses a heparanase (*hpse2*) (**Figure 4E**), which breaks down heparan sulfate proteoglycans (HSPs). HSPs are essential for fin growth^48^. Because heparanases liberate and release FGF from the HSP bond, into the extracellular matrix, the distal mesenchyme’s specific expression of HSPs may indicate this cell type also acts to amplify the effects of FGF signaling from the AER in a feedforward loop.

Finally, we identified an unknown third cell type that uniquely expresses *col4a1, chl1b* (**Figure S4B**) and the thrombospondin extracellular matrix glycoprotein *comp* (**Figure 4F**, right), also known as *thbs5*^54^. Based on its location in the center of the pectoral fin at 72 hpf and expression of thrombospondin gene, we initially labeled these unknown cells as the ‘central cells’ (**Figure 4B**). To validate that all of these terminal cell types derived from the *tbx5a*+ mesoderm, we used CRISPR to edit the *tbx5a* locus. As expected from the lineage map, all four markers of the terminal cell types disappeared (**Figure S4D**).

### Three-dimensional spatial model of pectoral fin identifies distal mesenchyme and tenocytes

The pectoral fin is a complex structure that initially grows out from a bud on the flank of the embryo (approximately 100μm in length)^55^. Gene expression boundaries are difficult to distinguish using traditional enzyme-based *in situ* hybridization (ISH). To understand the location of each of the cell types identified using our transcriptional atlas, we used confocal microscopy together with Hybridization Chain Reaction (HCR) to build a three-dimensional map of the expression of key marker genes with respect to one another, in the emerging fin at 48 hpf (**Figure 5A**). 3D-imaging of each of the top gene markers revealed that each cell type occupied a distinct spatial niche in the fin. At the center of the bud is the *emilin3a* cartilage condensate cells, which are flanked by *myod* expressing muscle cells that migrate in from the somites^56^. Surrounding these are *tbx5a*+, *fgf10a*+ mesodermal cells that will contribute to the growth of the fin bud. At the most distal end of the mesoderm are the *robo3*+ cells, that lie underneath the AER, marked by the expression of *prdm1a*, confirming our identification of these cells as separate from the AER, in a region identified cytologically as distal mesenchyme^55^ (**Figure 5B-D**).

**Figure 5.**
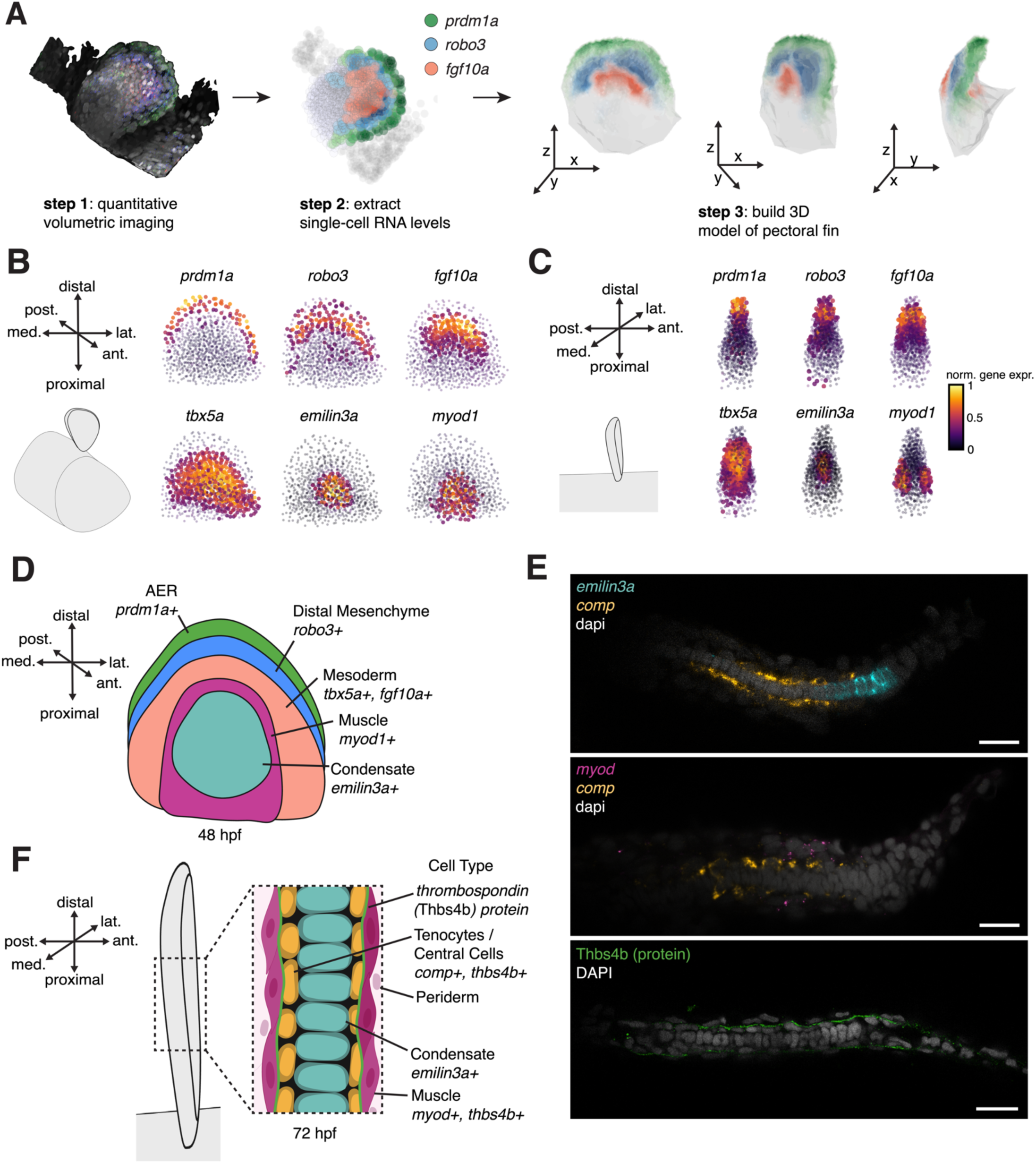
3D imaging and modeling reveals the cell type organization of the pectoral fin bud. A) Schematic diagram of the design of the 3D modeling pipeline for visualizing spatial gene expression in a common 3D fin space. Front B) and side view C) of the gene expression patterns in our 3D common pectoral fin space. post. posterior; ant. anterior; lat. lateral; med. medial. D) Summary 3D diagram of pectoral fin cell types based on gene expression in common fin space at 48 hpf. E) A representative cross-sectional view of 72 hpf wild-type pectoral fin labeled with fluorescent *in situ* hybridization (FISH), *emilin3a* in turquoise, *comp* in yellow (top), *myod1* in magenta, *comp* in yellow (middle) and IHC expression of Thbs4b in green (bottom). F) Summary 3D diagram of a cross-section of pectoral fin at 72 hpf, demonstrating tenocytes lie in between muscle and cartilage condensate and Thbs4b protein is expressed in between tenocytes and muscle. Scale bar is 20µm.

Because the expression level of the specific central cell marker *comp* was too low to be visualized by HCR, we used an alternative method that involves enzymatic deposition of a fluorescent substrate. In addition, because *comp* is only strongly expressed at 72 hpf, we imaged this marker at that time point. These results revealed that the central cells lie in a monolayer between the *emilin3a* expressing cartilage condensate and the *myod* expressing muscle on both the anterior and posterior sides of the pectoral fin (**Figure 5E**, top and middle). This localization, together with the expression, provided an important clue as to the role of these cells.

Thrombospondins play an essential role in the myotendinous junctions in the early zebrafish embryo^57,58^, with heteropentamers of *thbs4* and *thbs5* (*comp*) being secreted by tenocytes. Examination of *thbs4b* expression showed a pattern of expression similar to *comp*, with a speckled pattern across the 72 hpf fin (**Figure S4C**). Moreover, Thbs4b protein was observed in a thin layer between the muscle and tenocyte cells (**Figure 5E**, bottom). We therefore propose that the central cells are tenocytes, that produce *thbs4b* and *thbs5* (*comp)*, in order to attach the cartilage to the muscle (**Figure 5F**). In support of this, the central cells also express fibril-associated collagens with interrupted triple-helices (FACIT) collagens: *col22a1*, *col12a1a* and *col12a1b* (**Figure 4E**). These results are consistent with studies reporting the expression of the Scleraxis transcription factor gene (*scxa*) in the pectoral fin^59,60^. This new analysis of the cells deriving from the *tbx5a* progenitor population lays the groundwork for understanding the role of the different signaling factors in pectoral fin formation.

### Full model of signaling regulation in zebrafish pectoral fin mesoderm

Having constructed a 3D map of the pectoral fin, we next sought to characterize how the cell types within it communicate through each signaling pathway. When overlaid onto a map of the mesodermally-derived fin cell fates, the cell abundance changes enabled us to predict the role of each signaling factor on each pectoral fin cell type (**Figure 6A**). We then validated these predictions by visualizing expression of bud progenitors and bud mesoderm using WISH when the inhibitors were added at their earliest time point (**Figure 6B**), as well as the mesoderm and four differentiated cell fates when the inhibitors were added later at 24 hpf, after the bud mesoderm has already been established (**Figure 6C**). For cell types with well-characterized requirements for a given pathway, these experiments aligned with expectations. For example, differential cell type abundance and WISH validation recapitulated previous studies that FGF, Wnt and RA are all essential in the formation of the *tbx5a*+/*fgf24*+ fin bud progenitors and *tbx5a*+/*fgf10a*+ mesoderm as well as all the resulting cell types that originate from these cells^16–19^ (**Figure 6A-C, D**). Interestingly, Wnt inhibition results in a complete loss of the cleithrum marker, *runx2b*, while FGF inhibition leads to a strong reduction (**Figure 6C**). Since a recent study demonstrated that portions of the cleithrum originate from the cardio-pharyngeal mesoderm and neural crest^61^, this result shows that the *tbx5a* progenitors not only contribute to the cleithrum but non-autonomously play an important role in permitting the non-lateral plate cells to adopt the cleithrum fate.

**Figure 6.**
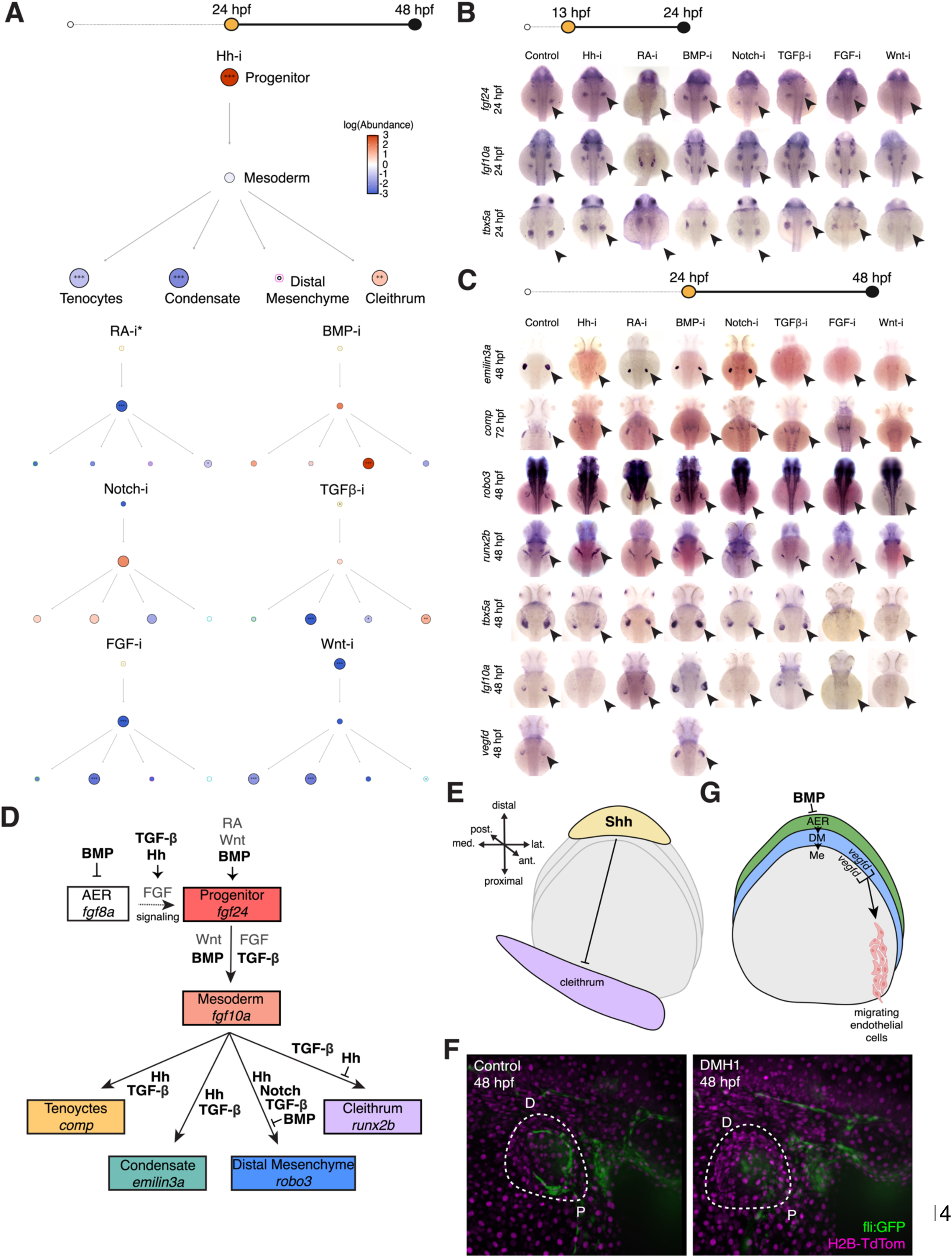
Identification of novel signaling pathway regulation in pectoral fin development. A) Platt-derived lineage-dependence graph, colored by significant (*, q < 0.05; **, q < 0.01; ***, q < 0.001) logFC in abundance of each cell type for each inhibitor compared to vehicle control added at 24 hpf and collected at 48 hpf, with exception of RA-i (*) added at 12 hpf because there was no effect on the pectoral fin when the inhibitor was added at 24 hpf. Dots are sized by the negative log of the q value, so more significant changes are larger. B) WISH of early pectoral fin bud mesoderm-specific genes (*fgf24*, *fgf10a, tbx5a*) in all perturbations and controls at 24 hpf. All inhibitors and vehicle controls added at 13 hpf. Black arrow indicates pectoral fin bud. C) WISH of terminal pectoral fin cell type-specific genes (cartilage condensate, *emilin3a*; tenocytes, *comp*; distal mesenchyme, *robo3*; cleithrum, *runx2b*) in all perturbations and controls at 48 hpf. All inhibitors and vehicle controls added at 24 hpf. Black arrowhead indicates pectoral fin region. D) Diagram depicting known (grey) and novel (bolded black) signaling pathway regulation of the pectoral fin mesoderm lineage. Dotted line indicates signaling from AER to bud progenitors. E) Diagram of Shh’s regulation of proximal and distal pectoral fin cell types. F) *Fli*-GFP embryos treated with DMH1 or DMSO control at 24 hpf and collected at 48 hpf. White dashed circle indicates pectoral fin, D, distal; P, proximal. G) Diagram of BMP’s non-autonomous regulation of endothelial cell migration.

In contrast to Wnt, FGF and RA inhibition that caused a loss of the fin bud, inhibition of the other four pathways had specific and different effects. Inhibition of Notch, for example, caused a strong reduction or loss of the *robo3*+ distal mesenchyme (**Figure 6C,D**). Whereas previous studies demonstrated that Hh is essential for formation of the pectoral fin^30,31^, we specifically found that Hh positively regulates the differentiation of the fin mesoderm into three of the terminal cell fates, cartilage condensate, tenocytes and distal mesenchyme (**Figure 6A-C**). Hh continued to regulate the condensate even when the inhibitor was added up to 32 hpf (**Figure S5A**). Strikingly, inhibition of Hh signaling caused an increase in the level of cleithrum cells in the sequencing data (**Figure 6A)** and that was validated with the cleithrum marker *runx2b* (**Figure 6C**), as well as additional cleithrum markers *cd81b* and *col27a1a* (**Figure S5B**). While *shha* eventually ends up on one side of the fin bud, during the early stages of fin bud emergence *shha* is at the distal tip of the bud, close to most of the mesodermal derivatives but distant from the cleithrum (**Figure S5C**). This fits with a gradient model whereby *shha* positively activates the mesodermal derivatives close to the signal but represses the most distal derivative, the cleithrum (**Figure 6D,E**).

### Pectoral fin mesoderm differentiation requires TGF-β

Examining TGF-β inhibition revealed that it did not affect the mesoderm at early stages since *fgf24*, *fgf10a* and *tbx5a* expression was normal at 24 hpf when the TGF-β inhibitor was added at the earliest time point, 13 hpf (**Figure 6B**). However, when the TGF-β inhibitor was added at 24 hpf, after the bud mesoderm had formed, all of the terminal mesodermal derivatives were reduced or absent at 48 hpf (**Figure 6A,C**). We noticed that TGF-β inhibition had similarities to FGF inhibition, including a partial inhibition of the cleithrum marker *runx2b* (**Figure 6C**). While *fgf10a* expression in the mesoderm was unaffected by TGF-β inhibition (**Figure 6C**), examination of the FGF ligands expressed in the AER revealed that *fgf8a* (**Figure S5D**) and *fgf4* (**Figure S5E**) were absent. We also examined the expression of the Shh ligands, *shha* and *shhb*, and found that neither were expressed in the absence of TGF-β signaling (**Figure S5E**). As Shh signaling was previously shown to regulate expression of FGF in the AER^30,31^, we propose that TGF-β regulates mesodermal differentiation indirectly via regulating *shha* and *shhb* expression. However, TGF-β inhibition also causes a *reduction* in cleithrum cell type abundance (*runx2b*, **Figure 6C**), which is the opposite of the increase caused by loss of Hh. Therefore we propose that TGF-β also acts to induce the cleithrum cell fate (**Figure 6D**).

### BMP indirectly regulates endothelial migration through the distal fin mesenchyme

A previous study showed that upregulation of BMP signaling using morpholino knockdown of a miRNA that targets the BMP inhibitor Noggin3 causes defects in the pectoral fin including a reduction of the cartilage condensate and a loss of the cleithrum^62^. In contrast, we see that inhibition of BMP signaling causes no change in the cleithrum, with a partial reduction in cartilage as indicated by the markers *runx2b* and *emilin3a*, respectively (**Figure 6A,C**). Intriguingly, BMP inhibition causes an increase in the amount of AER as shown with *prdm1a* and *fgf8a* (**Figure S5D**) and distal mesenchyme as shown with *robo3* expression (**Figure 6A,C**). Given that the AER is directly adjacent to the distal mesenchyme, it is likely that BMP’s negative regulation of distal mesenchyme may be indirectly mediated by the AER, which forms first. Because of the axon guidance properties of the Robo proteins, we looked for alterations in neuron distribution when Bmp was inhibited, but did not find any obvious changes (data not shown). However, we also found that two vascular endothelial growth factor signal genes (*vegfd, vegfaa)*, which work as chemoattractants for blood vessels, are expressed in the pectoral fin^63,64^ (**Figure S5F)**. While the levels of *vegfaa* as observed in WISH are very low in the pectoral fin and in many embryos undetectable (data not shown), in agreement with our sequencing results, *vegfd* is clearly expressed in the distal mesenchyme and distal region of the core mesoderm (**Figure 6C, Figure S5F**). Similar to *robo3*, BMP inhibition causes a strong increase in *vegfd* expression, with the *vegfd* no longer confined to the most distal tip of the fin (**Figure 6C**). We therefore examined the blood vessels in the pectoral fin using a transgenic line that expresses GFP in the vessels^65^. While the blood vessels in control fins migrate to the distal portion of the fin, when BMP is inhibited, the vessels fail to migrate into the fin (**Figure 6F**), clearly demonstrating a role for BMP signaling in pectoral fin vascularization. We propose that when the chemoattractant *vegfd* is not restricted to the distal tip, the blood vessels no longer receive a clear signal to migrate out to the end of the pectoral fin (**Figure 6D,G**).

## Discussion

Here, we have measured the consequences of interrupting each of seven major signaling pathways on the transcriptomes of each cell type in the developing zebrafish embryo. We applied embryo-scale single-nucleus RNA-seq to hundreds of individual zebrafish embryos treated with different pathway inhibitors, sequencing over 2 million cells in total. We measured how disabling signaling at different stages of development changes cell type proportions and gene expression within each cell type. From these effects, we have constructed a map describing which cell types depend on which pathways and at what stages of embryonic development.

This new resource will enable investigators to quickly determine whether any of the major signaling pathways influence their cell type or tissue of interest, providing a foundation for more in-depth investigation. Additionally, it will facilitate the identification of gene expression changes in this cell type or tissue when signaling is altered.

In order to construct these maps, we had to develop new tools that statistically quantify the effects of such perturbations on cell type proportions and gene expression. To do so, we developed Hooke and Platt, described in a companion paper^12^ (Duran *et al.* see related manuscript file). That paper describes the statistical principles, algorithms and software that powers Hooke and Platt and demonstrates their utility for interpreting genetic perturbations. Here, we use Hooke and Platt to not only define which cell types depend on which signals and how, but to watch how effects of chemical perturbations propagate through the cell lineage. This enables us to draw inferences about which cell types depend directly on signaling as opposed to indirectly through their lineage ancestors. Together, the two studies demonstrate that Hooke and Platt can operate over a diverse range of single-cell perturbation experimental designs to draw causal inferences about the control of embryonic development.

By applying our new tools to our single-cell atlas of chemically perturbed embryos, we observed changes in the proportions and transcriptomes of the overwhelming majority of cell types in the atlas, including in very rare cell types. Inhibiting each pathway induces a distinct, consistent pattern of changes to cell type proportions, reflecting the pervasive use of the major pathways across the vertebrate embryo. Gene expression analysis showed that inhibiting the pathways induced reproducible and consistent modulation of classic pathway targets and effectors in many cell types across the embryo, as well as responses from genes not known to be targets, including lineage-determining transcription factors. Although many cell types were depleted or lost when a given pathway was inhibited, others displayed only transcriptional phenotypes. In some cases, such as forebrain neural progenitors that fail to fully express *notch3* in Notch-i embryos, transcriptional phenotypes in ancestral cells anticipated changes to the abundance of their descendants in the lineage. Further mining of such patterns with more sophisticated tools designed to reconstruct gene regulatory networks could expand our maps of how each cell type interprets signals to make fate decisions and specialize its broader program of gene regulation.

Although this resource captured effects of modulating signaling across the embryo, we focused on the pectoral fin in detail to explore the potential of our approach for guiding deeper study. The pectoral fin was directly impacted by the inhibition of all seven of the pathways, including pathways not previously known to play a role in fin or limb development. We used our tools to map lineage relationships in the pectoral fin, mapping out how the mesodermally derived cells, including two previously uncharacterized populations, descend from a common ancestor. Detailed analysis of marker genes using several imaging techniques enabled us to construct a 3D model of the fin, which in turn served as a basis for a deeper exploration of how signaling regulates its development. Specifically, we describe previously uncharacterized control of pectoral fin development by TGF-β and BMP signaling. We found that TGF-β is required for proper expression of *fgf8a* in the AER, and that inhibiting TGF-β interrupts the critical feedforward signaling loop between the bud mesoderm and the AER that drives fin outgrowth. We also found that TGF-β is important later in the differentiation of the mesoderm into the four terminal cell types. Additionally, we showed that BMP governs fin blood vessel patterning indirectly by regulating the size and distribution of the distal mesenchyme and the most distal region of the mesoderm, which both express *vegfd*, a key chemoattractant for blood vessels. Our “deep dive” into the signaling requirements of the pectoral fin not only confirmed that our approach is sound, it also revealed important new and likely conserved features of pair appendage development. It is not surprising that the pectoral fin uses all the signaling pathways - it is an organ that is mechanically intricate, evolutionarily ancient, constructed from cells of multiple cell lineages, and undergoes complex morphogenetic transitions throughout embryonic and post-embryonic development. However, our study underscores the value of perturbing not just one pathway but all of them, and then monitoring those perturbations on each cell type, for dissecting the interplay of signaling in complex developing tissues.

These data constitute the first comprehensive, embryo scale, genome wide analysis of signaling requirements of vertebrate embryonic cell types. They therefore constitute a new and substantial contribution to the growing single-cell atlas of how different perturbations alter the development of the zebrafish embryo. We distilled these data into a map of signaling requirements for each cell type that, as we demonstrate by exploring the pectoral fin, can guide further experiments into the organization of tissues, how cell types within them signal to one another, and how those interactions govern cell fate. The map captures signaling requirements of diverse cell types, a substantial number of which have been previously reported, underscoring the quality of the data and the power of our approach. However, it also includes many previously unreported requirements. Our focused follow-up experiments dissecting the novel requirements in the pectoral fin validate and suggest the resource has great potential for discoveries in other organs and tissues. More broadly, our approach of using single-cell transcriptomic phenotyping to study chemically perturbed zebrafish embryos, which allows one to target pathways that are difficult to study with a genetic approach, is a powerful means of defining how cell types interpret signals in the developmental program.

Our single-cell atlas of chemically perturbed embryos, together with new software packages Hooke and Platt will serve as a foundation for further experiments to dissect the mechanisms that direct cell fate decisions and tissue morphogenesis. For example, although we documented many cell types and genes that depend on each of the signals we interrupted, the specific components that transduce and potentiate those signals in each cell type are yet to be defined. Layering chemical perturbations onto mutants and then performing single-cell sequencing could shed light on the kinases and transcription factors that each cell type uses to interpret signals. Moreover, although our analysis of gene expression changes uncovered evidence of cross-talk between pathways, how exactly the pathways are interacting with one another, and which downstream pathway components are used in each cell type, is not clear. These are important questions because although signals pattern the embryo, generating distinct cell fates in different anatomic positions, cells at similar positions (and receiving the same signals) often make different choices, and thus interpret those signals differently^2^. Defining the particular receptors and pathway components needed for cells to commit to each fate is critical for discriminating whether signals license a cell’s potential fates or merely commit it to one of them.

## Methods

### KEY RESOURCES TABLE

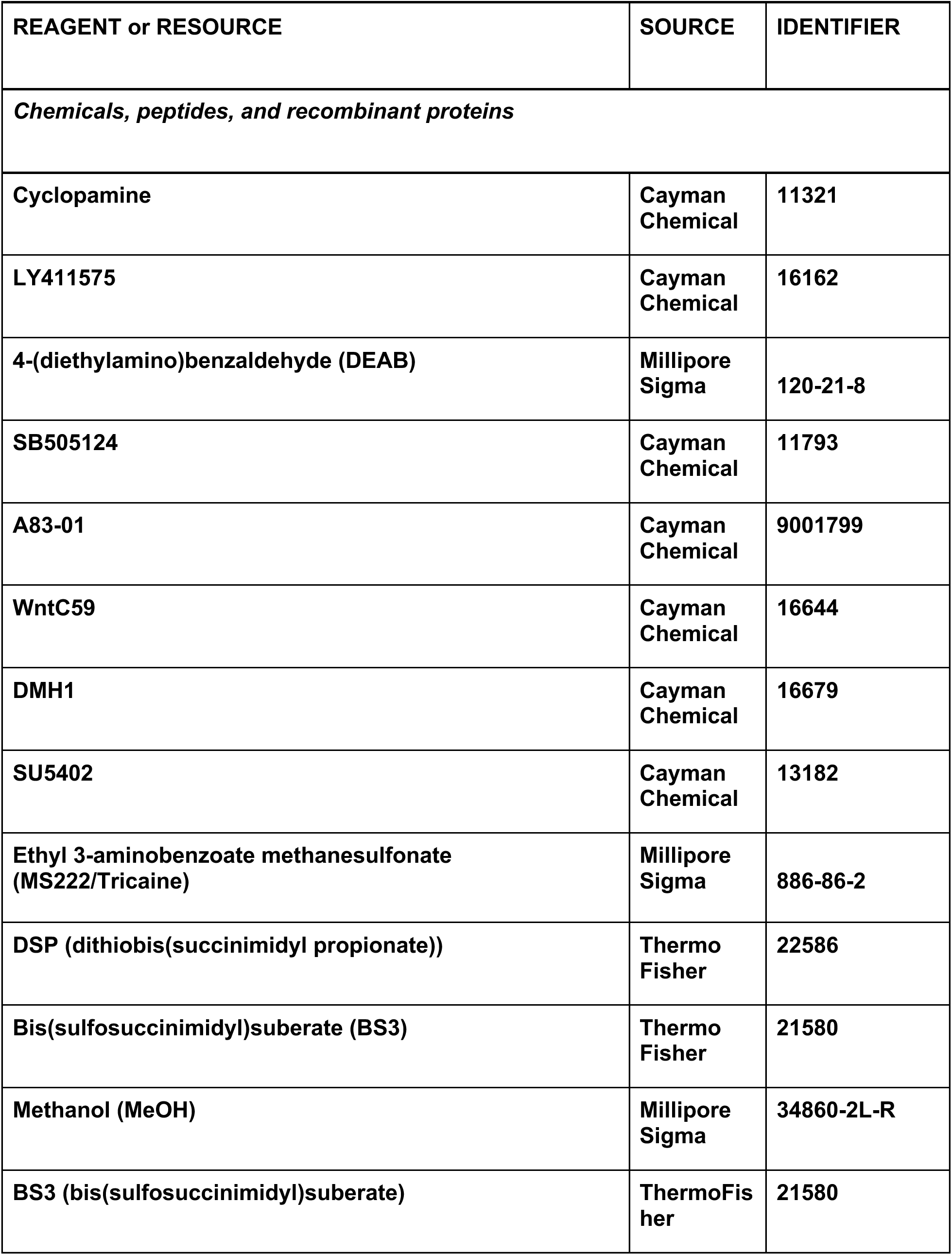

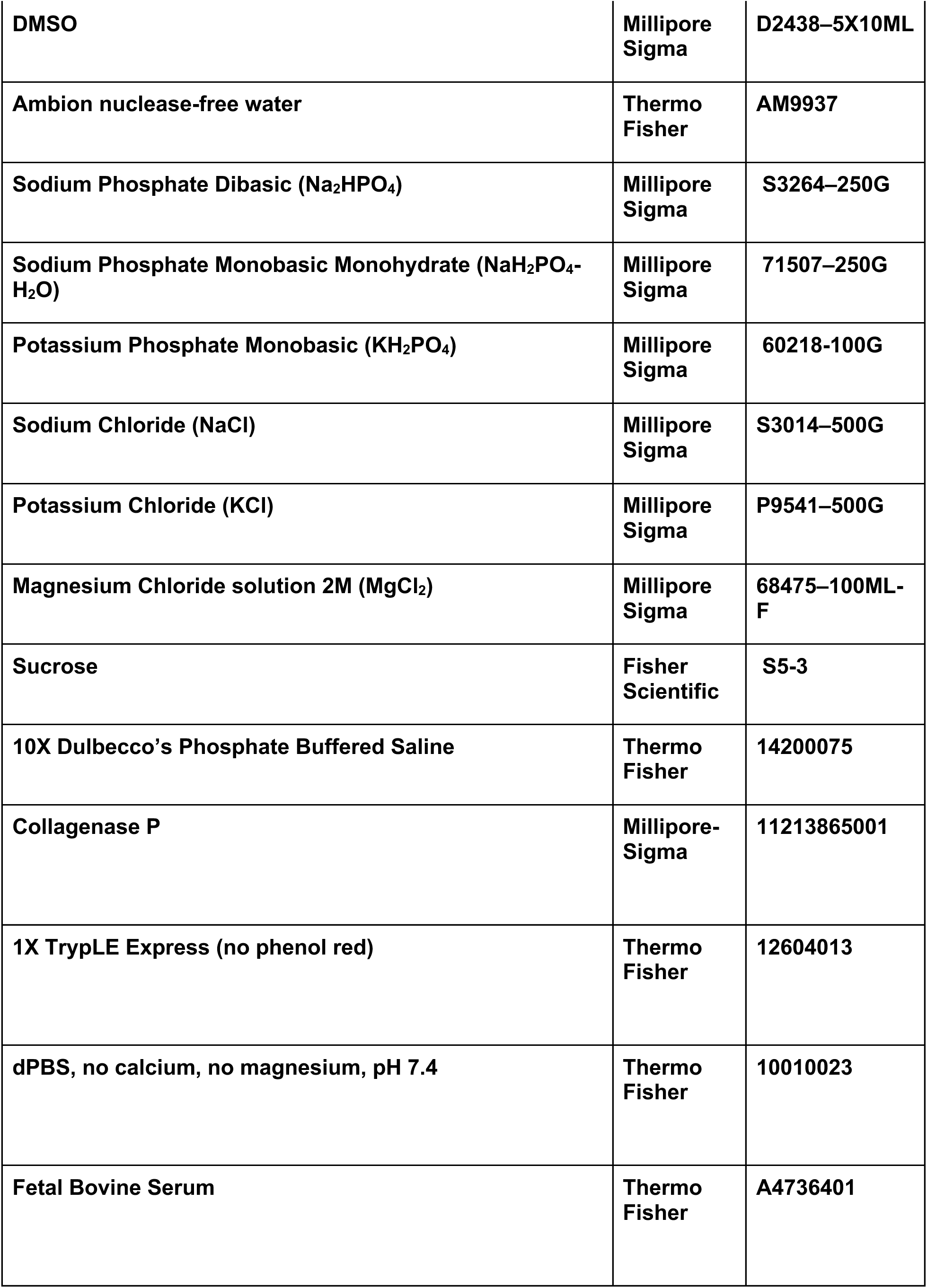

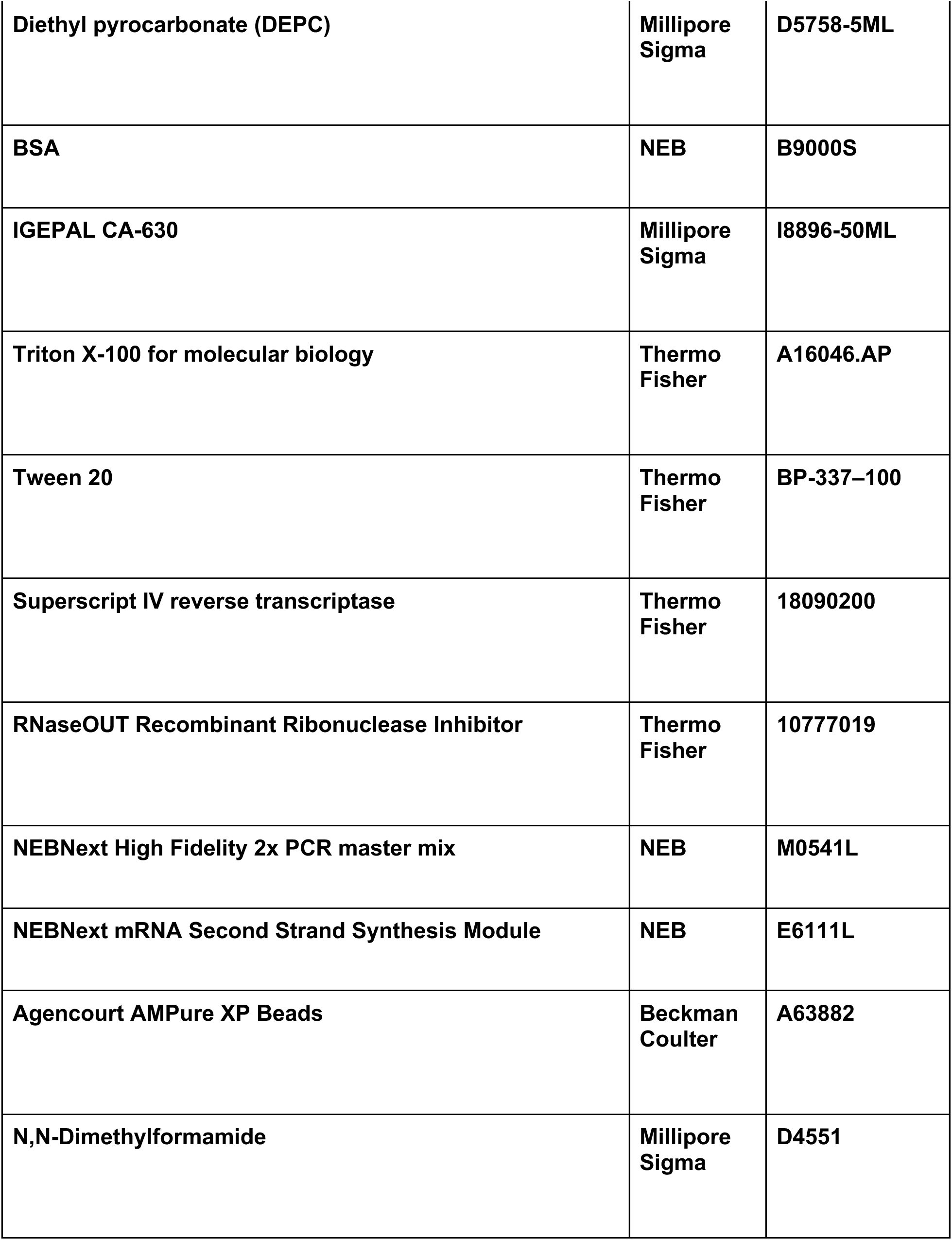

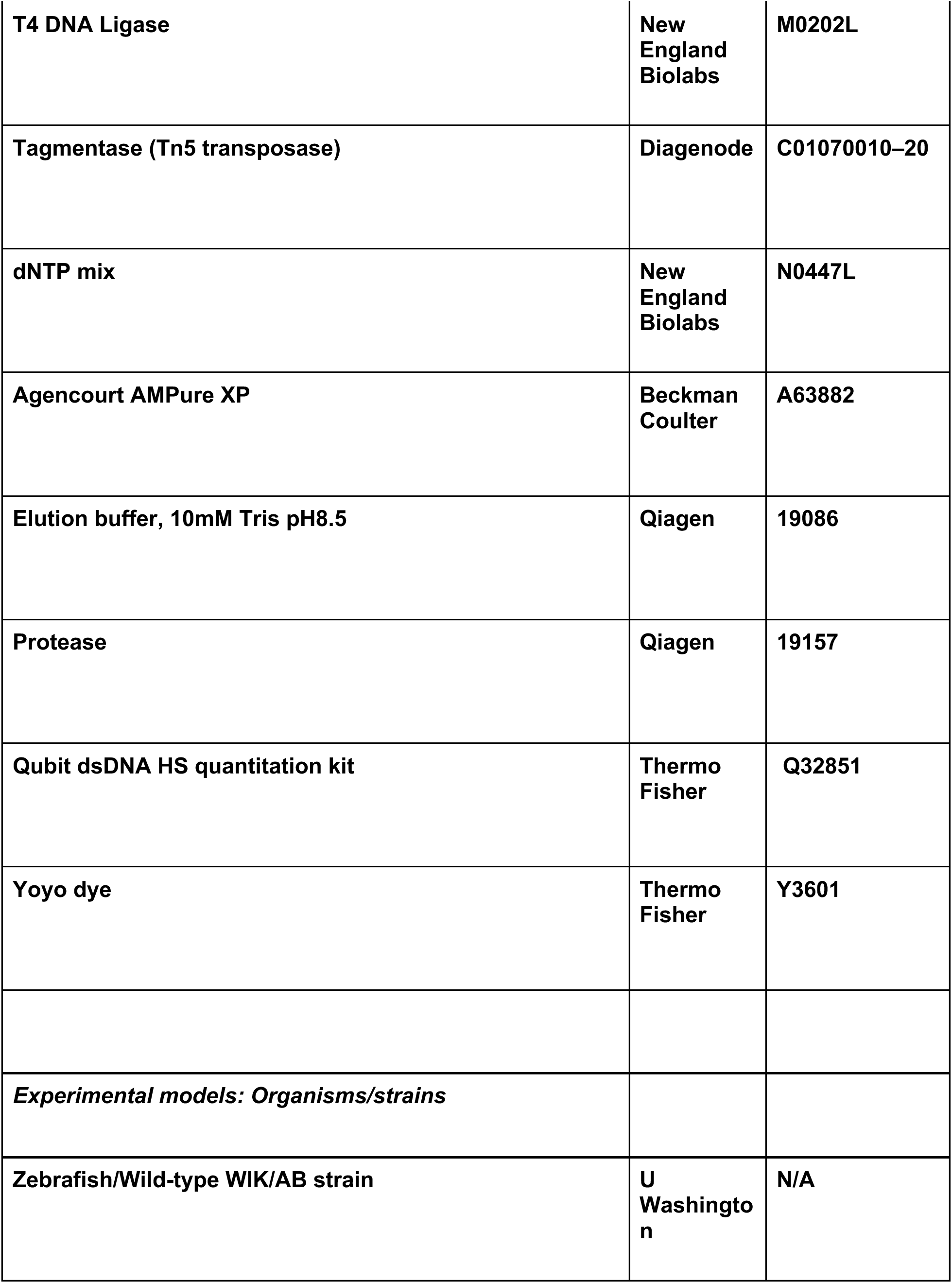

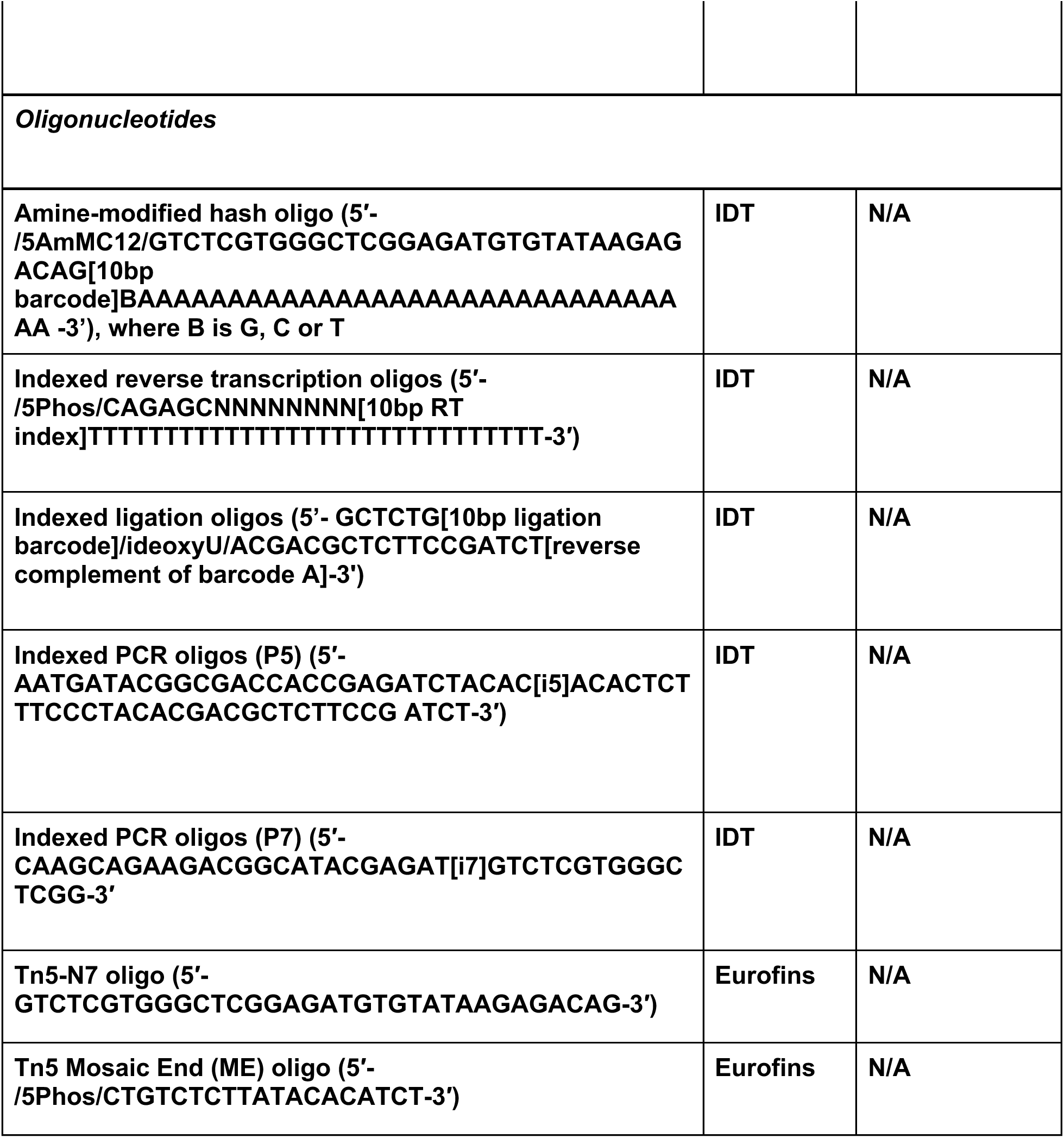

### Animal rearing, staging and stocks

Staging followed^6^ and fish were maintained at 28.5°C under 14:10 light:dark cycles. Fish stocks used were wild-type AB or WIK/AB except for the use of the *fli:GFP* line^65^ for imaging the pectoral fin vasculature. Fish were anesthetized prior to imaging or dissociation with MS222 and euthanized by overdose of MS222. All procedures involving live animals followed federal, state and local guidelines for humane treatment and protocols approved by Institutional Animal Care and Use Committees (protocol #4405-02) of the University of Washington.

### Chemical Inhibitor Screen

Embryos were exposed to the inhibitors 100µM Cyclopamine (Cayman, cat. no. 11321),10µM DEAB (Millipore Sigma, cat. no. 120-21-8) or 10µM LY411575 (Cayman, cat. no. 16162) at shield stage with vehicle controls 1% Ethanol for Cyclopamine control or 0.24% DMSO for all other inhibitor controls. Embryos were exposed to the inhibitors 50µM SB505124 (Cayman, cat. no. 11793), 50µM A83-01 (Cayman, cat. no. 9001799), 20µM WntC59 (Cayman, cat. no. 16644), 40µM DMH1 (Cayman, cat. no. 16679) or 20µM SU5402 (Cayman, cat. no. 13182) at 8-somite stage with vehicle control 0.24% DMSO. Embryo media was replaced and inhibitors or vehicle were replenished every 24 hours until time of collection.

### Preparation of Barcoded Nuclei

Individual zebrafish embryos (36, 48 and 72 hpf) were manually dechorionated with forceps and transferred to a 10cm petri dish containing 1X TrypLE (Thermo Fisher, cat. no. 12604013) and MS222 (Millipore Sigma, cat. no. 886-86-2). Embryos were dissociated into single cells according to the protocol described in Saunders *et al.*, 2023. For round 1, cell lysis and fixation followed the protocol described in Martin *et al.* (2023), with an additional 5μl of C6 amine-modified hash DNA oligo (10uM, IDT, 5′-/5AmMC12/GTCTCGTGGGCTCGGAGATGTGTATAAGAGACAG[10bp barcode]BAAAAAAAAAAAAAAAAAAAAAAAAAAAAAAAA −3’) mixed into the Hypotonic Lysis Buffer Solution B^66^. For round 2, the same protocol was followed except the fixative Bis(sulfosuccinimidyl)suberate (BS3) (75μl 100mM BS3 with 5mL MeOH) was used instead of DSP (dithiobis(succinimidyl propionate)).

### sci-RNA-seq3 library construction

The fixed and hashed nuclei were processed according to the following protocol https://www.ncbi.nlm.nih.gov/pmc/articles/PMC9839601/pdf/nihms-1846803.pdf^66^.

### Sequencing, read processing, and cell filtering

Sequencing, read processing and cell filtering were performed according to Saunders *et al.*, 2023. An enrichment cutoff of 2.5 was set based on the distribution of enrichment ratios (Figure S1C). Pipelines for generating count matrices from sci-RNA-seq3 sequencing data are available at https://github.com/bbi-lab/bbi-dmux and https://github.com/bbi-lab/bbi-sci.

### Single cell RNA-seq analysis

After RNA and hash-quality filtering, data were processed using the Monocle3 (develop branch, v.1.3.1) workflow using defaults except where specified: *align_cds(residual_model_formula_str* = *‘∼log10(n.umi)’)*, *reduce_dimension(max_components* = *3, preprocess_method* = *‘Aligned’)* and *cluster_cells* where k was scaled by the number of cells.

### Cell projection and label transfer

Projection and label transfer were performed similarly to Saunders *et al.* (2023), using zscapetools, with some exceptions that are detailed in Duran *et al.* (see related manuscript file)^12^. The PCA rotation matrix, batch-correction linear model, and UMAP transformation were computed and saved during the processing of the reference dataset. This computation was done on two levels: first, with all combined reference cells (global reference space), and then, in each of the subgroups (sub-reference space). Projection group labels were transferred using the majority label of its annotated nearest neighbors (*k* = 10) using annoy, a fast, approximate nearest-neighbor algorithm (https://github.com/spotify/annoy, v.0.0.20). The query dataset was split into these subgroups and query subgroups were projected into their respective sub spaces. Cell type labels were again transferred using the majority vote of reference neighbors (*k* = 10).

### Differential cell abundance testing

Abundance testing was computed using a poisson log normal model with Hooke^12^ (Duran *et al*., see related manuscript file) as described in the user guide (https://cole-trapnell-lab.github.io/hooke/). After cell type annotation, counts per cell type were summarized per embryo, generating an embryo × cell type matrix. The embryo × cell type matrix was stored as a Hooke cell_count_set object. Counts were compared across chemical perturbations and their paired controls. Log abundances were estimated for each condition at each timepoint and the fold change is computed between perturbations and controls along with standard errors under the Poisson log normal model. P-values were calculated using a Wald test and corrected with Benjamini-Hochberg multiple testing correction.

### Differential gene expression analysis

Differential gene expression across perturbations was computed for every cell type using *Platt*^12^ (Duran *et al*., see related manuscript file). Briefly, *Platt* models each gene’s expression in each cell type, in each treatment condition using a generalized linear regression model (GLM). The magnitude of a perturbation’s effect on a gene in a cell type was estimated by subtracting the coefficients for the paired control from that of the perturbation, which yields an estimate of the log fold-change in expression. *ashr*^67^ was imposed to regularize effect estimates due to cell types with low counts.

### In situ hybridization

Colorimetric in situ hybridization used digoxygenin labeled probes using standard conditions^68^. HCR probes were designed by Molecular Instruments, Inc. and were used following the manufacturer’s protocol (https://files.molecularinstruments.com/MI-Protocol-RNAFISH-Zebrafish-Rev10.pdf). Tyramide signal amplification was performed according to the Lauter et al. (2011) protocol.^69^

### Cartilage staining

Alcian blue staining followed an online procedure (SDB Online Short Course, Zebrafish Alcian Blue, https://www.sdbonline.org/sites/2008ShortCourse/Zebrafish_AlcianBlue.pdf), with the exception that embryos were raised in 1-phenyl-2-thiourea to suppress pigment formation instead of bleaching.

### Immunohistochemistry

A standard immunohistochemistry protocol was performed^70^.

### Imaging

Embryos live-imaged were anesthetized with MS222 and photographed on a Nikon AZ100 microscope. For colorimetric RNA *in situs* and alcian blue stained samples, embryos were put into 70% glycerol and imaged on a Nikon AZ100 microscope. For the alcian blue samples, most of the yolk was manually removed and the sample was placed under a coverslip. For confocal imaging of HCR, TSA and IHC, a Zeiss LSM 900 upright confocal microscope equipped with either a plan-apochromat 20x/1.0 or 10x/0.5 water objective was used to image the whole mount pectoral fin samples. The larvae were mounted dorsal side down in 1.2% low-melt agarose in 1X PBS. After the molten agarose solidified, the sample was inverted dorsal side up and remounted in 60 mm petri dishes for imaging. Images were corrected for color balance and display levels as necessary with all conditions within each analysis grouping corrected identically.

### Confocal Imaging Analysis

For analysis, the raw confocal image data was deconvoluted using Zen Blue. The DAPI channel was then used to perform 3D nucleus segmentation using a custom-trained CellPose model^71^. For each of the twelve confocal images, this produced 3D masks indicating the boundary for each nucleus within the image volume. Custom Python software was written to specifically identify nuclei located within the pectoral fin. These pectoral fin nuclei formed the basis of all subsequent gene expression analyses. In addition to the DAPI channel, each imaging dataset contains three additional channels containing fluorescence measurements for HCR probes targeting mRNA corresponding to three genes of interest. To quantify expression levels for each gene, we calculated the average intensity within the corresponding fluorescence channel across all pixels contained within each nucleus mask. Each single-nucleus estimate was then averaged with its five nearest spatial neighbors to reduce the influence of technical and biological noise. This process led to three-dimensional maps of gene expression within the pectoral fin for each of our twelve imaging datasets. Custom Python software was written to combine the twelve datasets into an average spatial atlas of pectoral fin gene expression. The key to this process was co-projecting each dataset into a shared three-dimensional reference space. This was done in three main steps: (i) standardizing fin pose and orientation, (ii) calculating an average pectoral fin shape across the twelve datasets, and (iii) using non-rigid registration to align each pectoral fin with the average fin. After this process, nuclei from each dataset were located within a shared reference space defined by the average fin. K-means clustering was then used to create 1000 ‘average cells’, each representing 12-13 actual nuclei from across the twelve imaging datasets. Gene expression levels for each average cell were obtained by averaging the levels across its constituent nuclei. Only nuclei from experiments that included a given gene were used to calculate its average expression.

### gRNAs

Two gRNAs directed against the first exon of the *tbx5a* gene (GTATGTAGTCTGCGATGACG and TACAGGCCTCTCCGACATGGCGG) were mixed with Cas9 (IDT) and injected into one-cell embryos.

### Hierarchical annotation and subclustering of pectoral fin

To annotate pectoral fin mesoderm cell types in the reference atlas (Saunders *et al*., 2023), the subclustered mesoderm was subdivided, embedded in three dimensions with UMAP and subclustered based on *tbx5a* and *runx2b* gene expression. Clusters within pectoral fin were then assigned annotation based on the expression of marker genes (Hooke new_cell_count_set & pseudobulk_ccs_for_states), and monocle3 top_markers function, significance determined by a two-sided likelihood test with multiple testing correction, based on literature using anatomical-term gene lists from ZFIN database^72^. The full revised zebrafish reference atlas with these new pectoral fin annotations is published in Duran *et al*. (see related manuscript file)^12^.

### Pectoral fin lineage construction

Pectoral fin lineage was constructed using Platt from wildtype time series data in the reference atlas that spanned 18 hpf to 96 hpf (Duran et *al*., see related manuscript file)^12^.

## Data Availability

The accession number for the single cell RNA-seq data reported will be available soon. Processed data files used in the paper analysis are available for download at https://cole-trapnell-lab.github.io/chemfish/ under CC-BY-NC.

## Project Acknowledgments

We thank the Brotman Baty Institute Advanced Technology Lab for support with sequencing and the data processing pipeline. We thank Dr. Sanjay Srivatsan, Dr. Dave Raible, Dr. Doug Fowler, Dr. Alice Berger, Dr. Edith Wang, Dr. Jay Shendure and Beth Martin for helpful discussions as we were developing this project.

## Author Contributions

E.B. and D.K. conceived the project. E.B. and D.K. designed experiments. L.S. trained E.B. on dissociation and nuclei collections. E.B., D.K., H.L., B.H., A.T. and L.S. did dissociation and nuclei collections. E.B., D.K., H.L., B.H., A.T. and L.S. performed sci-RNA-seq3 experiments. D.K. performed all micro-injections. D.K. performed in situ hybridizations with E.B. E.B. and M.D. performed computational analyses. E.B. annotated pectoral fin cell types. N.L. and P.A. performed imaging experiments. N.L. constructed 3D pectoral fin model. E.B. D.K. and C.T. wrote the manuscript with input from all co-authors. C.T. and D.K. supervised the project.

## Supplemental Information

**Supplemental Table 1** – DACT by perturbation, timepoint of drug addition and timepoint of collection.

**Supplemental Table 2** – Pectoral Fin DEGs.

## Supplemental Figures

**Supplemental Figure 1.**
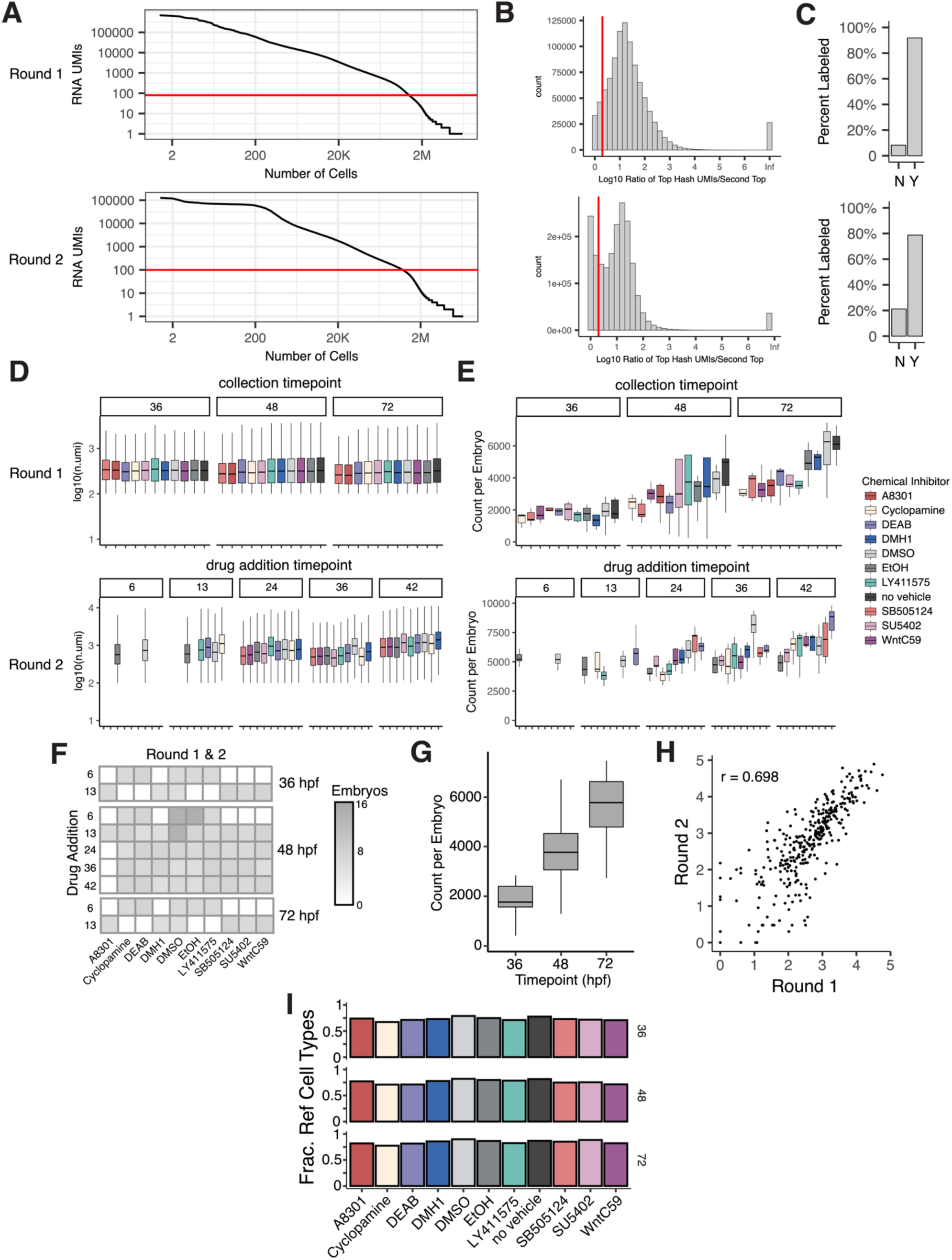
Experimental metrics for inhibitor experiments. A) Ranked plot of the number of unique molecular identifiers per cell (UMIs). Barcodes with UMIs above the red line (80 UMIs in round 1, 100 UMIs in round 2) were considered cells. B) Distribution of the highest UMI hash barcode per cell divided by the second highest UMI hash barcode per cell. Cells with a ratio of 2.5 or greater and total hash UMIs > 5 after subtracting background hash UMI level, were considered a high quality assignment and were included in our study. C) Percentage of cells uniquely labeled with a ratio >= 2.5 and total hash UMIs >= 5 (Y) with a ratio < 2.5 and total hash UMIs < 5 (N). D) Boxplot of the number of mRNA UMIs per cell across all embryos per perturbation (color) and collection time point in round 1 (top) or inhibitor addition time point in round 2 (bottom). E) Boxplot of the number of cells per embryo collected across all embryos per perturbation (color) and collection time point in round 1 (top) or inhibitor addition time point in round 2 (bottom). All inhibitors were added at one time in round 1 (6 or 13 hpf depending on inhibitor) and all embryos were collected at one time in round 2 (48 hpf). F) Heatmap of the number of embryos collected at each inhibitor addition and collection time point for each perturbation. G) Boxplot of the number of cells per embryo captured across controls collected at all three time points in round 1. H) Scatterplot of the percent of the control embryos made up by each cell type in the 48 hpf collection in round 1 vs round 2. These results demonstrate there is high reproducibility in cell type abundance across the two rounds. I) Fraction of the number of unique cell types found in our experiments out of total possible cell types found in matching timepoints in the reference atlas for each perturbation and collection time point in round 1. In both rounds, ethanol was the vehicle for Cyclopamine and DMSO was the vehicle for all other inhibitors.

**Supplemental Figure 2.**
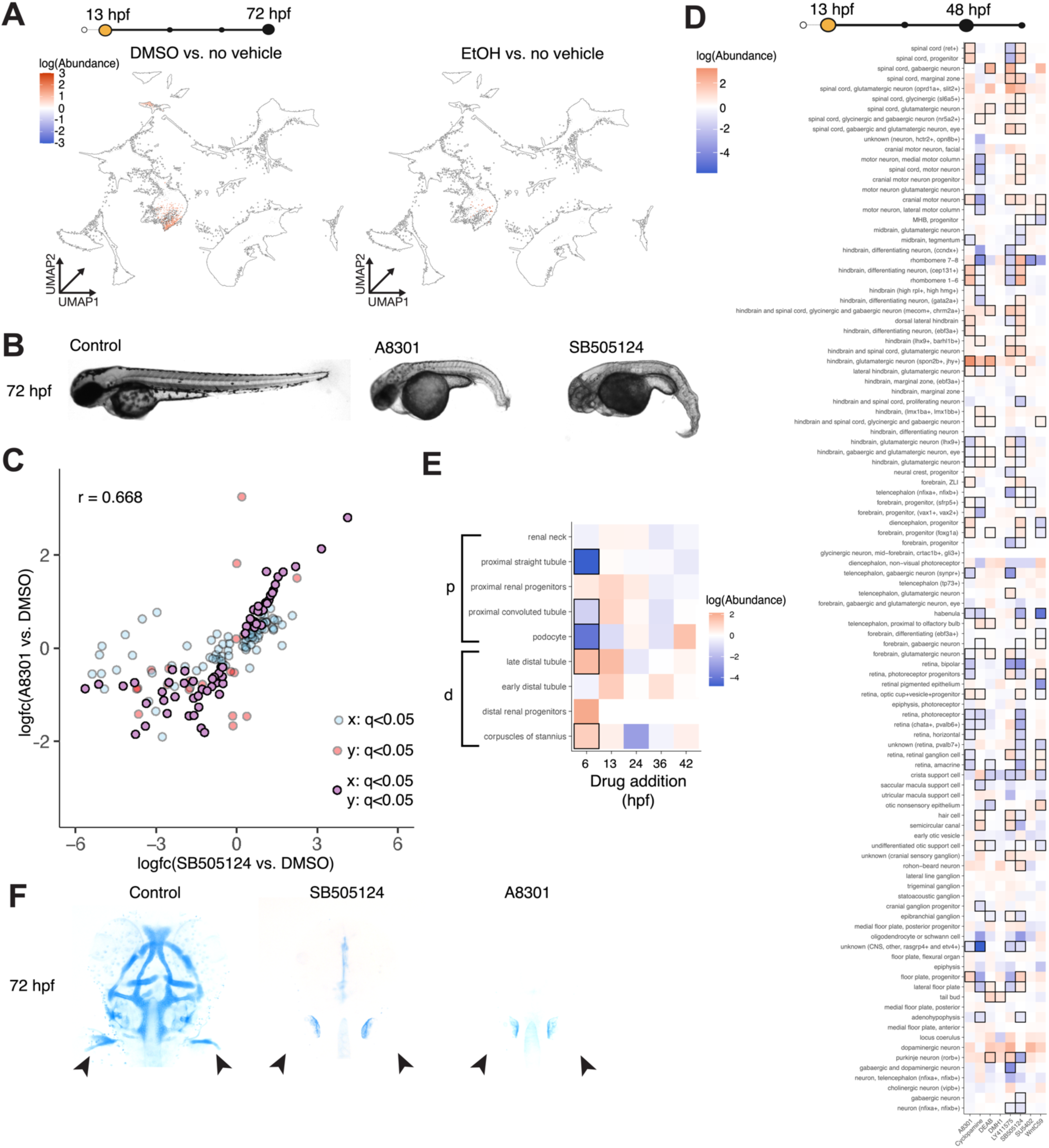
Controls for abundance phenotyping of chemically inhibited embryos. A) UMAP embedding, colored by significant (q<0.05, PLN regression) logFC in abundance of each cell type for DMSO vehicle control (left) and ethanol control (right) vs. no vehicle control at 48 hpf, vehicle controls added at 13 hpf. These results demonstrate that the vehicle controls (DMSO and EtOH) do not have a significant effect on cell type abundance. B) Images of embryos treated with each of the two TGF-β inhibitors, SB505124 and A8301 and DMSO vehicle control at 72 hpf. C) Scatterplot of logFC in abundance of each cell type for each of the two TGF-β inhibitors, SB505124 and A8301 added at 13 hpf, across all collection time points. Cell types significant in both SB505124 and A8301 are colored in purple and outlined in black, cell types significant in only SB505124 or A8301 are colored in blue or red, respectively. D) Heatmap of all CNS cell types, ordered by projection groups: spinal cord, motor neurons, midbrain, ear, hindbrain, eye, forebrain, cranial sensory ganglia, CNS other and colored by logFC in abundance of each cell type in 13 hpf addition of each inhibitor vs vehicle control, collected at 48 hpf. Black boxes indicate significance (q<0.05, PLN regression). E) Heatmap of pronephros cell types, colored by logFC in abundance of each cell type across each time point of DEAB (RA-i) addition vs vehicle control, collected at 48 hpf. Black boxes indicate significance (q<0.05, PLN regression). F) Images of alcian blue stained 72 hpf embryos treated with each of the two TGF-β inhibitors, SB505124 and A8301 and DMSO vehicle control at 13 hpf. Black arrow indicates pectoral fin.

**Supplemental Figure 3.**
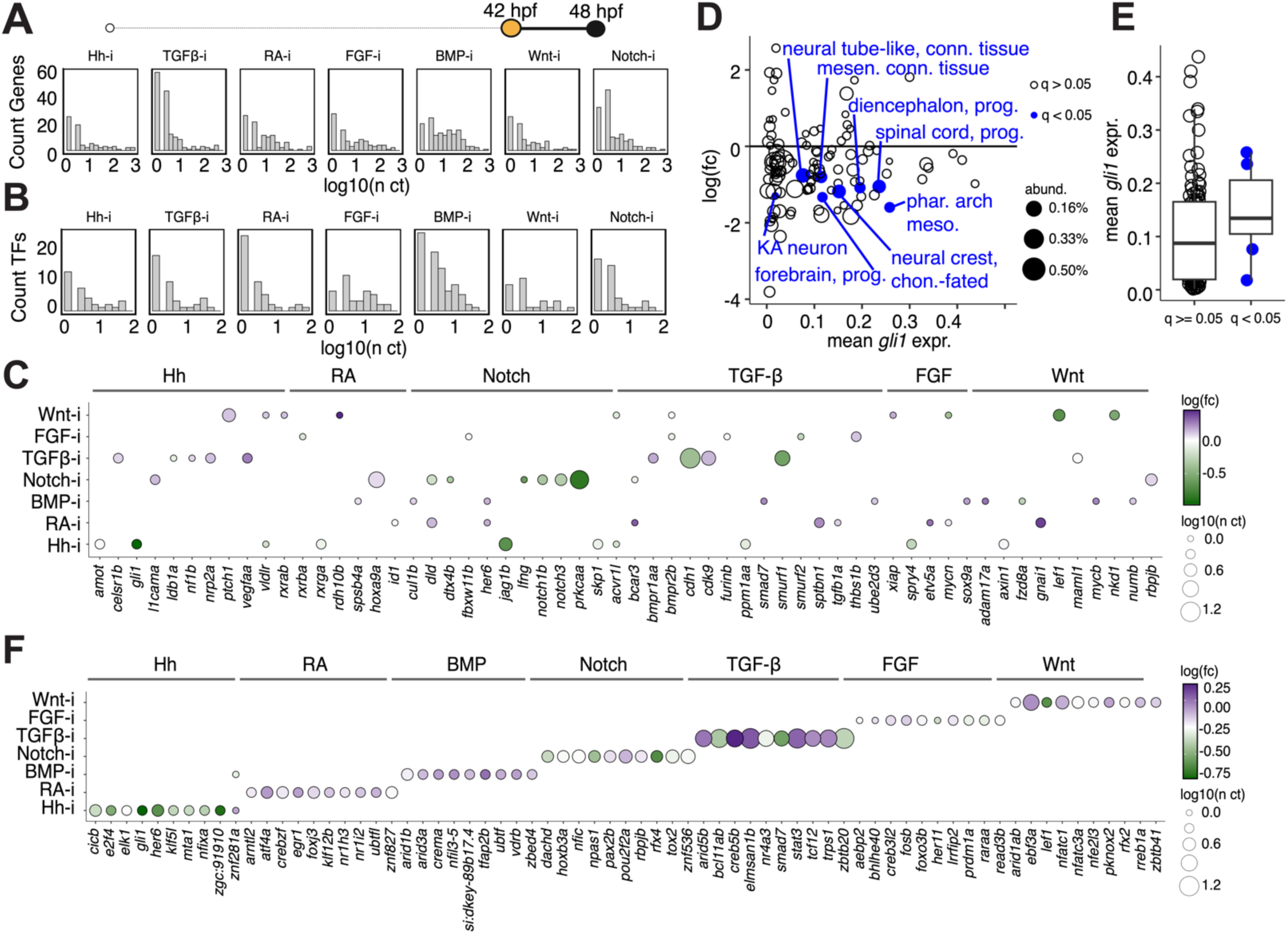
Summary statistics of differentially expressed genes. A) Distribution of the number of cell types (n ct) that each gene is significantly (q<0.05) differentially expressed in. B) Distribution of the number of cell types that each TF is significantly differentially expressed in. C, F) Dotplot heatmap of the averaged effects of fold changes in (C) expected downstream genes and (F) top ten most commonly differentially expressed TFs for each signaling pathway across all cell types. The ten most highly expressed downstream genes in the reference atlas are plotted. Dots are sized by the number of cell types with a significant logFC for each gene and perturbation. D) Scatterplot of the mean *gli1* expression of control cells in the reference atlas versus logFC in cell type abundance in cyclopamine-treated, Hh-i embryos. Significant DACTs are labeled and colored blue. Dots are sized by the percent abundance of the cell type in the reference atlas. E) Boxplot of mean *gli1* expression for non-significant DACTs (q >= 0.05) vs. significant DACTs (q < 0.05). All data shown in this figure are from 42 hpf inhibitor addition and 48 hpf collection, except the y-axis of panel D is from the 6 hpf addition of cyclopamine.

**Supplemental Figure 4.**
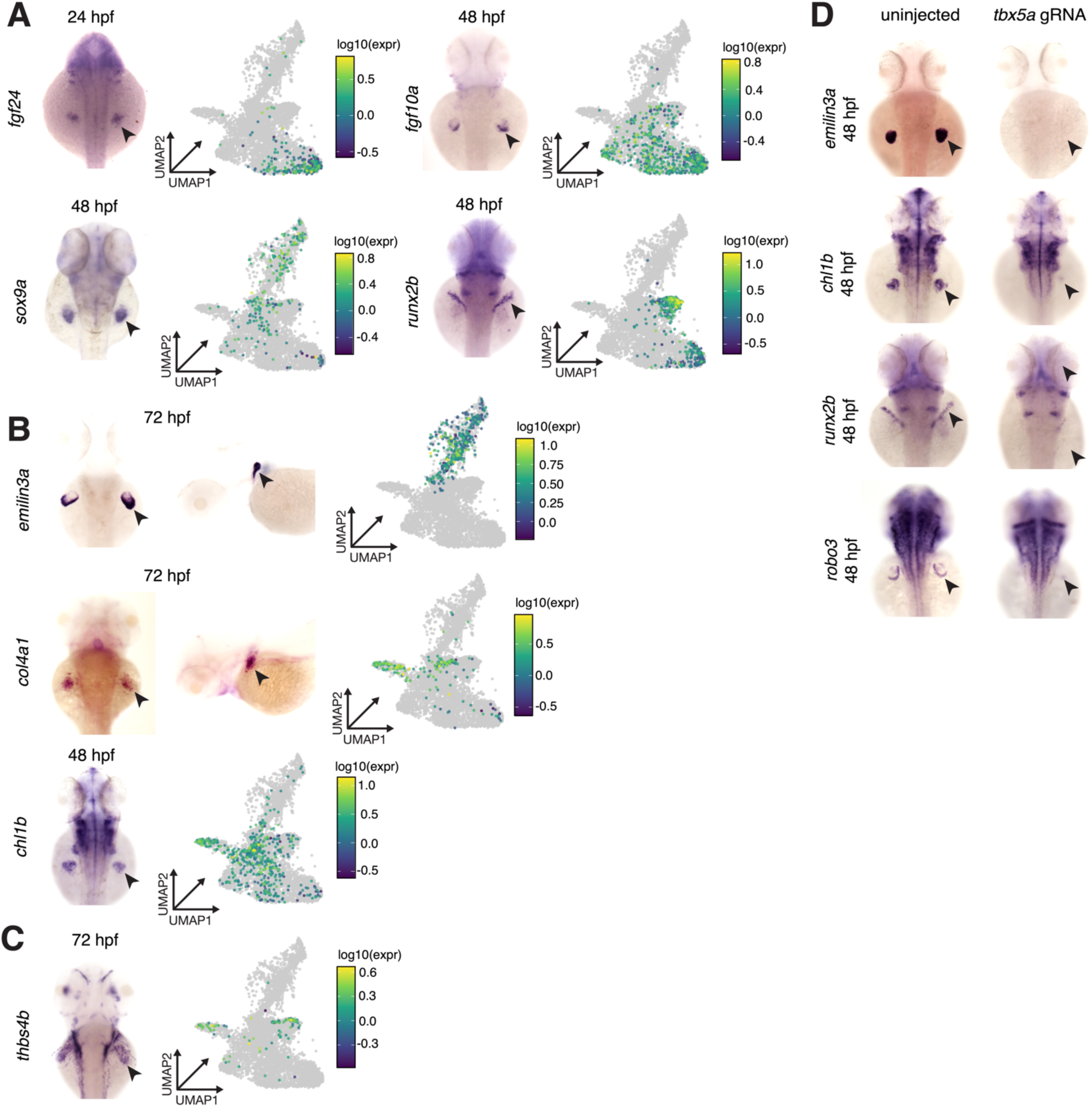
Extended top marker expression of pectoral fin mesoderm cell types. A) WISH of canonical pectoral fin cell type-specific genes in control embryos and UMAP colored by log10 of gene expression (right). B) WISH of pectoral fin cell type-specific genes, *emilin3a* (condensate), *col4a1* (central cells) and *chl1b* (central cells) (left) and UMAP colored by log10 of gene expression (right). C) WISH of central cell marker gene *thbs4b* in control embryo at 72 hpf (left) and UMAP colored by log10 of gene expression. Thbs4b is also expressed in cleithrum. All time points are shown in all UMAPs. D) WISH of terminal pectoral fin cell type-specific genes (cartilage condensate, *emilin3a*; central cells, *chl1b*; distal mesenchyme, *robo3*; cleithrum, *runx2b*) in *tbx5a* F0 crispants at 48 hpf. Black arrowhead indicates pectoral fin region.

**Supplemental Figure 5.**
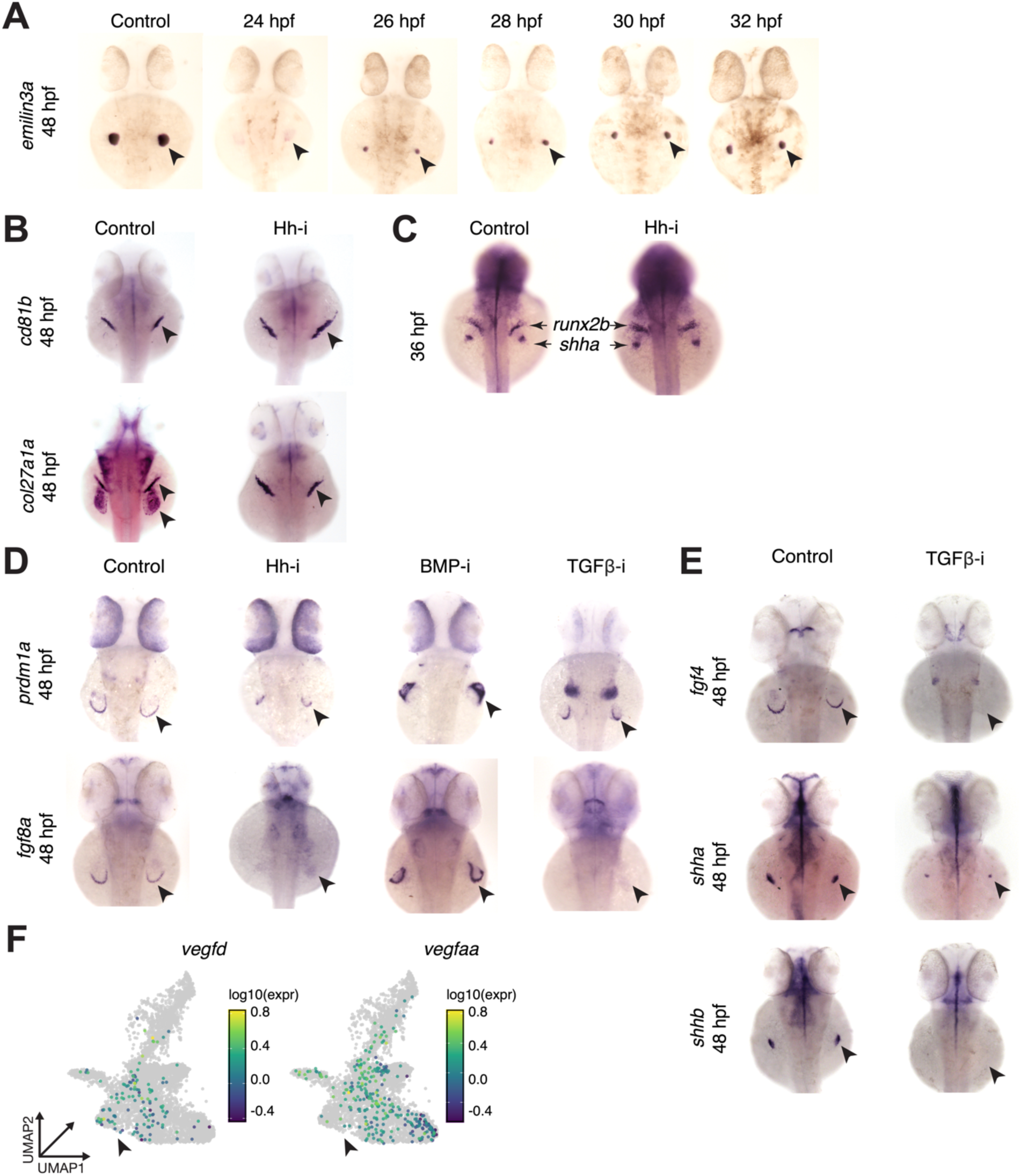
Extended validation of proposed signaling pathway regulation in pectoral fin mesoderm. A) WISH of control and cyclopamine-treated, Hhi, embryos stained for cartilage condensate-specific genes (*emilin3a*). Cyclopamine was first added at the indicated time point. B) WISH of cleithrum-specific genes (*cd81b*, *col27a1a*) in cyclopamine-treated, Hh-i embryos at 48 hpf. Chemical inhibitor added at 24 hpf. C) Dual WISH of cleithrum (*runx2b*) and *shha* expression in Hh-i, cyclopamine-treated and vehicle control at 36 hpf. Chemical inhibitor added at 6 hpf. D) WISH of AER marker genes, *prdm1a and fgf8a* in a panel of multiple chemically inhibited embryos at 48 hpf. Inhibitor added at 24 hpf. SB505124 was used to inhibit TGF-β unless otherwise noted. E) WISH of *fgf4* and shh (*shha, shhb*) expression in TGFβ-i, SB505124-treated and vehicle control at 48 hpf. Chemical inhibitor added at 24 hpf. Black arrow indicates pectoral fin. F) Pectoral fin UMAP colored by log10 expression of *smoc1* gene expression. Black arrow indicates distal mesenchyme-fated mesoderm.

